# Phylogenomics reveals patterns of ancient hybridization and differential diversification contributing to phylogenetic conflict in *Populus* L. and *Salix* L

**DOI:** 10.1101/2023.01.04.522772

**Authors:** Brian J. Sanderson, Diksha Ghambir, Guanqiao Feng, Nan Hu, Quentin C. Cronk, Diana M. Percy, Francisco Molina Freaner, Matthew G. Johnson, Lawrence B. Smart, Ken Keefover-Ring, Tongming Yin, Tao Ma, Stephen P. DiFazio, Jianquan Liu, Matthew S. Olson

**Affiliations:** Department of Biological Sciences, Texas Tech University, Lubbock, TX 79409-3131 USA; Department of Botany, University of British Columbia, Vancouver, BC, V6T 1Z4 Canada; Universidad Nacional Automoa de Mexico, Hermosilla, Mexico; Horticulture Section, School of Integrative Plant Science, Cornell University, Cornell AgriTech, Geneva, New York 14456 USA; Departments of Botany and Geography, University of Wisconsin-Madison, Madison, WI 53706, USA; Key Laboratory of Tree Genetics and Biotechnology of Jiangsu Province and Education Department of China, Nanjing Forestry University, Nanjing, China; Key Laboratory of Bio-Resource and Eco-Environment of Ministry of Education & College of Life Sciences, Sichuan University, Chengdu 610065, China; Department of Biology, West Virginia University, Morgantown, WV, 26506 USA; State Key Laboratory of Grassland Agro-Ecosystem, Institute of Innovation Ecology & College of Life Sciences, Lanzhou University, Lanzhou 730000, China

**Author notes:** These authors contributed equally to this work (Brian J. Sanderson, Matthew S. Olson). The Jackson Laboratory for Genomic Medicine, 10 Discovery Drive, Farmington, CT 06032 USA.

**Keywords:** Salicaceae, hybridization, sequence capture, ASTRAL species tree, concatenated tree

## Abstract

Despite the economic, ecological, and scientific importance of the genera *Populus* L. (poplars, cottonwoods, and aspens) and *Salix* L. Salicaceae (willows), we know little about the sources of differences in species diversity between the genera and of the phylogenetic conflict that often confounds estimating phylogenetic trees. *Salix* subgenera and sections, in particular, have been difficult to classify, with one recent attempt termed a ‘spectacular failure’ due to a speculated radiation of the subgenera *Vetrix* and *Chamaetia*. Here we use targeted sequence capture to understand the evolutionary history of this portion of the Salicaceae plant family. Our phylogenetic hypothesis was based on 787 gene regions and identified extensive phylogenetic conflict among genes. Our analysis supported some previously described subgeneric relationships and confirmed polyphyly of others. Using an f_branch_ analysis we identified several cases of hybridization in deep branches of the phylogeny, which likely contributed to discordance among gene trees. In addition, we identified a rapid increase in diversification rate near the origination of the *Vetrix-Chamaetia* clade in *Salix*. This region of the tree coincided with several nodes that lacked strong statistical support, indicating a possible increase in incomplete lineage sorting due to rapid diversification. The extraordinary level of both recent and ancient hybridization in both *Populus* and *Salix* have played important roles in the diversification and diversity in these two genera.

**Supplementary data files will be provided by request to**

---

*This study is dedicated to the memory of George W. Argus (1929-2022) whose lifelong pursuit of understanding diversity in Salix laid the foundation for future salicologists*.

## Introduction

As methods to assess the congruence among the genealogical histories of genes across species have matured (Degnan et al. 2009; Young et al. 2020), the curious association between phylogenetic conflict and rapid diversification has suggested a link between population genetic and macroevolutionary processes (Parins-Fukuchi et al. 2021). Although most genomic regions are expected to reflect the speciation and diversification history of a taxonomic group (species tree), two primary factors contribute to conflict between gene genealogies and species history. Incomplete lineage sorting (ILS) results from the persistence of polymorphism across multiple diversification events and the subsequent random fixation of polymorphism among different lineages (Wu 1991). The influence of ILS is particularly strong during periods of rapid speciation, when effective population sizes are large and when long-term balancing selection results in persistence of polymorphisms (Edwards 2009; Pease et al. 2013; Wang et al. 2020). Second, interspecific gene flow due to hybridization has the potential to generate discordance among high proportions of gene trees (McVay et al. 2017; Morales-Briones et al. 2020). Unlike ILS, however, the patterns of gene discordance are biased with over-representation of one hybrid topology (Green et al. 2010; Durand et al. 2011; Patterson et al. 2012). When taxonomic groups have hybridized throughout diversification, the effects of hybridization on gene tree conflict can span multiple species within a clade (Malinsky et al. 2018). The impacts of hybridization may be particularly common in plants, where fertility of interspecific crosses may be maintained well after speciation (Grant 1981).

Contemporary populations of poplars (*Populus*) and willows (*Salix*) are widely known to hybridize, with important evolutionary and ecological consequences along hybrid zones (Brunsfeld et al. 1992; Hardig et al. 2000; Schweitzer et al. 2004; Lexer et al. 2010; Chhatre et al. 2018; Wang et al. 2020). The impacts of hybridization on the evolution and diversification are evident in both genera, where multiple independent chloroplast capture events have occurred early in their diversification (Smith et al. 1990; Brunsfeld et al. 1992; Liu et al. 2017; Wang et al. 2020; Gulyaev et al. 2022). This history of hybridization creates challenges for the reconstruction of phylogenetic histories of poplars and willows (Percy et al. 2014). Recent progress using genome-wide data sets, however, have constructed well-supported taxonomic relationships within both genera (Wagner et al. 2020; Wang et al. 2020; Wagner et al. 2021a; Gulyaev et al. 2022; Wang et al. 2022), but the sources of conflict among gene trees have not been fully investigated, especially in *Salix*.

Poplars and willows are integral components of temperate, boreal, and arctic ecosystems throughout the northern hemisphere and many species have significant cultural, medical, and economic importance (Stettler et al. 1996; Argus 1997). Phytochemical diversity in these genera spans an impressive array of secondary metabolites, including the aspirin and its derivatives (Desborough et al. 2017) and defensive chemicals such as phenolic glycosides, condensed tannins and hydroxycinnamate derivatives (Tsai et al. 2006; Philippe et al. 2007; Boeckler et al. 2011; Keefover-Ring et al. 2022). Morphological variation in both genera ranges from dwarf creeping *Salix* in alpine and arctic zones that were once categorized as a separate genus (Stettler et al. 1996; Argus 1997) to large *Populus* trees in subtropical zones. Many more species are recognized in *Salix* (approx. 450-520 species; Argus 2010) than in *Populus* (approx. 100 species; Shu 1999), suggesting that either *Salix* began to diversify much earlier than *Populus* or speciation rates have increased in *Salix*.

*Populus* and *Salix* are the two largest genera in the Salicaceae and all but one species across both genera are dioecious (Rohwer et al. 1984). The Salicaceae whole genome duplication unites all genera except *Azara*, which lacks the duplication (Cronk et al. 2015). Within the Salicaceae, *Populus* and *Salix* are united by a striking synapomorphy of flowers organized into aments or catkins (Meeuse 1975; Argus 2010; Eckenwalder 2010; Cronk et al. 2015) with seed dispersal via wind. This differs from their closest relative *Idesia*, which produces fleshy animal-dispersed fruits. *Populus* and *Salix* differ in that pollen is dispersed by wind in *Populus* and by insects or by both insects and wind in *Salix* (Sacchi et al. 1988; Tamura et al. 2000; Karrenberg et al. 2002), suggesting that factors underlying pollination mode may drive differences in the diversification rate between the two genera (Friedman et al. 2009; Wessinger 2021). The reduced floral structures in both genera exhibit relatively low variability and are used to discriminate among species only at the broadest taxonomic levels (Eckenwalder 1996; Argus 1997). Thus, plant stature and growth form, leaf morphology, and bud characteristics have been important characters for species identification (Dorn 1976; Argus 2010; Eckenwalder 2010) despite the high intraspecific variability and propensity for plasticity of these traits (Wu et al. 2015).

Here we seek to understand the sources of phylogenetic conflict in both *Populus* and *Salix* by comparing their historical patterns of hybridization and relative rates of diversification, both factors that can contribute to conflict with species trees. Our ASTRAL phylogeny was based on genome-wide sequence capture loci from a large set of mostly North American and Asian diploid *Salix* and *Populus* species. The phylogeny of a subset of these *Salix* samples was previously investigated using DNA barcode markers, but the resulting tree lacked resolution (Percy et al. 2014). Also, a phylogeny of the majority of the *Populus* samples were previously analyzed by Wang et al. (Wang et al. 2020) using a different set of whole genome loci, providing a “positive control” to confirm that our targeted sequence capture array designed for Salicaceae (Sanderson et al. 2020) successfully reconstructs the species tree. We use this tree to compare the impact of hybridization on gene tree discordance and the chronology and rates of diversification across these two genera. The specific goals of our study are to: 1) provide an integrated phylogenetic hypothesis of the sister genera *Populus* and *Salix*, 2) to estimate the timing and rates of diversifications of major clades within each genus, and 3) to assess the association between hybridization history and regions of the phylogeny that exhibit conflict among gene trees. Finally, we discuss the implications of our results in the context of other well-known hybridizing groups of species (syngameons) species such as oaks (Cannon et al. 2020) and pines (Buck et al. 2022).

## Methods

The 166 samples included in this study were drawn from new collections, older dried herbarium samples, and previously sequenced genomes, and represented all five *Populus* sections, all five *Salix* subgenera, and 25 *Salix* sections (Table S1). All species were considered diploids based on chromosome counts reported in www.tropicos.org except *S. discolor*, which is likely a polyploid, and *S. richardsonii*, for which there was no information concerning chromosome counts. *Salix* species were primarily native to North America, but also from Europe and Asia. Fourteen *Populus* samples (7 species) and 83 *Salix* samples (45 species) were genotyped using a custom sequence target capture kit designed for the Salicaceae (Supplemental Methods; Sanderson et al. 2020). An additional 6 outgroup species (*Azara dentata, A. integrifolia, A. lanceolata, A. microphylla, Carrierea calycina, Idesia polycarpa*, and *Poliothyrsis sinensis*), 54 poplar samples (26 species), and 8 *Salix* samples (7 species) were genotyped using whole genome sequencing. All sequences were assembled into putatively homologous gene sequences using the HybPiper pipeline (Johnson et al. 2016); for the whole genome sequences, this removed all loci except those included in the target capture array. After filtering, removal of alignments with paralogs, and removal of genes with excessively long branches in the gene tree, 787 alignments remained and were used for all downstream phylogenetic analyses (See Supplemental Methods for details; Table S2).

Two approaches were used for phylogeny estimation: 1) A two-step approach using ASTRAL that first estimated trees for each gene and then identified the best tree based on minimizing quartet distances among gene trees (Mirarab et al. 2014), and 2) identifying the maximum likelihood tree based on concatenating all genes in our sample. Full details of these analyses are provided in the Supplemental Methods. In brief for the two-step approach, gene trees were estimated using IQTREE 2.0.3 (Nguyen et al. 2015) and an ASTRAL tree of all individuals and the species tree (Rabiee et al. 2019) were inferred using ASTRAL-MP (v 5.12.2 (Yin et al. 2019). In our ASTRAL tree of all individuals, four species were represented by intraspecific samples that did not cluster together (Fig. S1): *Salix bebbiana, S. eriocephala, S. pseudomonticola*, and *Populus ningshanica;* these are represented by a postscript 1 & 2 in the figures and Table S1. For this reason, we used two individuals to represent each of these four species in the ASTRAL species tree. Both local posterior probability and 100 multilocus bootstraps were calculated for each ASTRAL tree. For the concatenation approach, a single alignment was constructed by concatenating all 787 genes (1,058,955 sites). IQTREE was used to estimate the most likely concatenated tree using the GTR+F+R10 substitution model. 1000 mulitlocus bootstraps and SH-alRT tests (Guindon et al. 2010) also were computed for this tree.

A dated species tree was calculated using *BEAST2 (Heled et al. 2010). Because of the long computation times required for sampling, five genes were chosen for dated tree estimation (9,264 sites: SapurV1A.0003s0350, SapurV1A.0045s0240, SapurV1A.0050s0650, SapurV1A.0139s0330, SapurV1A.0260s0050). These genes were first selected for high consistency with species tree topologies, minimized root-to-tip variance, and maximal tree length (Smith et al. 2018) and then screened for consistency of basal nodes with the ASTRAL species tree. Because these five genes represented <1% of the genes used for generating our species tree, we selected an additional five genes using the same criteria to assess the consistency of the results across different gene sets (8,568 sites: SapurV1A.0211s0160, SapurV1A.0789s0070, SapurV1A.0857s0020, SapurV1A.0900s0040, SapurV1A.1178s0060). See the Supplemental methods for additional information on gene selection and for *BEAST2 parameter settings. The calibration date for the root of the tree (divergence between *Azara* and all others) was drawn from a normal distribution with a mean of 65.0 Ma and standard deviation of 1.0 (following Wang et al. 2020), and the calibration of the crown clade of *Populus + Salix* was drawn from a normal distribution with a mean of 49 Ma and a standard deviation of 1.0 following Percy et al. (2014) and based on the *Pseudosalix handleyi* fossil (Boucher et al. 2003) from the Eocene Green River formation that has been dated at ca. 49 Ma (Smith et al. 2010). Using a wider standard deviation of 3.0 for the distributions resulted in much larger variance in the estimates of node ages, but only slight changes in the estimates for divergence times (Fig. S4).

Patterns of current and historical hybridization within the *Populus* and *Salix* clades were estimated using ABBA-BABA, f_4_, and f_branch_ analyses (Patterson et al. 2012; Malinsky et al. 2018) calculated using Dsuite (Malinsky et al. 2021). The f_branch_ analysis is heuristic and is designed to account for phylogenetic correlation among f_4_-ratio results calculated with phylogenetically correlated samples. The f_branch_ metrics assign significance to internal branches in the phylogeny when excess sharing of alleles that is consistent with hybridization is found across a clade (Malinsky et al. 2018; Malinsky et al. 2021). We generated separate VCF files for *Populus* and *Salix* to calculate f_4_-ratio and f_branch_ statistics using Dsuite using GATK v4.2.6.1 (McKenna et al. 2010) as detailed in the Supplemental Methods.

Two methods were used to address diversification rates across the Salicaceae. First, Bayesian Analysis of Macroevolutionary Mixtures (BAMM; Rabosky 2014) was used to identify credible shifts in the diversification rate across lineages with the *expectedNumberOfShifts* prior set to 1.0. Because our species-level sampling was not uniform across genera, we adjusted for non-random incomplete taxon sampling (Table S3). For each dated tree, two independent MCMC chains using different seeds were run for 10 million generations each resulting in ESS>400, and the 95% credible set of shift configurations was calculated after removing 10% for burn-in. Second, branch-specific diversification rates were estimated using the ClaDS model (Maliet et al. 2019) and calculated using data augmentation (Maliet et al. 2022). Finally, we used the “STructured Rate Permutations on Phylogenies” (STRAPP) test (Rabosky et al. 2015) to assess the association between categorical traits associated with wind versus insect+wind pollination, wind versus animal seed dispersal, and tree versus shrub-dwarf shrub growth form.

## Results

Of the 1216 loci that were targeted, 787 passed filtering criteria with an average of 162 individuals per gene (2.5% & 97.5% quantiles: 144 & 165), 1236 sites per sequence (2.5% & 97.5% quantiles: 673 & 2629), 305.0 parsimony-informative sites (2.5% & 97.5% quantiles: 134.7 & 673.7), 132.3 singletons (2.5% & 97.5% quantiles: 45 & 333.7), and 908.2 constant sites (2.5% & 97.5% quantiles: 452.0 & 1715.6) per gene. Our overall findings support monophyly for both *Salix* and *Populus*, and classification of the basal portions of each genus were consistent with prior morphological and molecular analyses (Figs. 1, S1; Eckenwalder 1996; Wagner et al. 2018; Wang et al. 2020; Gulyaev et al. 2022). Our phylogeny also supports recent discoveries of the polyphyly of the *Tacamahaca* and *Aigeiros* subgenera in *Populus* (Wang et al. 2020) and the polyphyly of the *Vetrix* and *Chamaetia* subgenera in *Salix*. Notably in *Salix*, our study and that of Wager et al. (2018) each identified at least 4 independent evolutionary events leading to dwarf willows (subgenus *Chamaetia*), which are prominent components of northern hemisphere arctic and alpine ecosystems. However, because these studies had little taxonomic overlap, it is difficult to discern whether we identified the same or different events.

**Figure 1.**
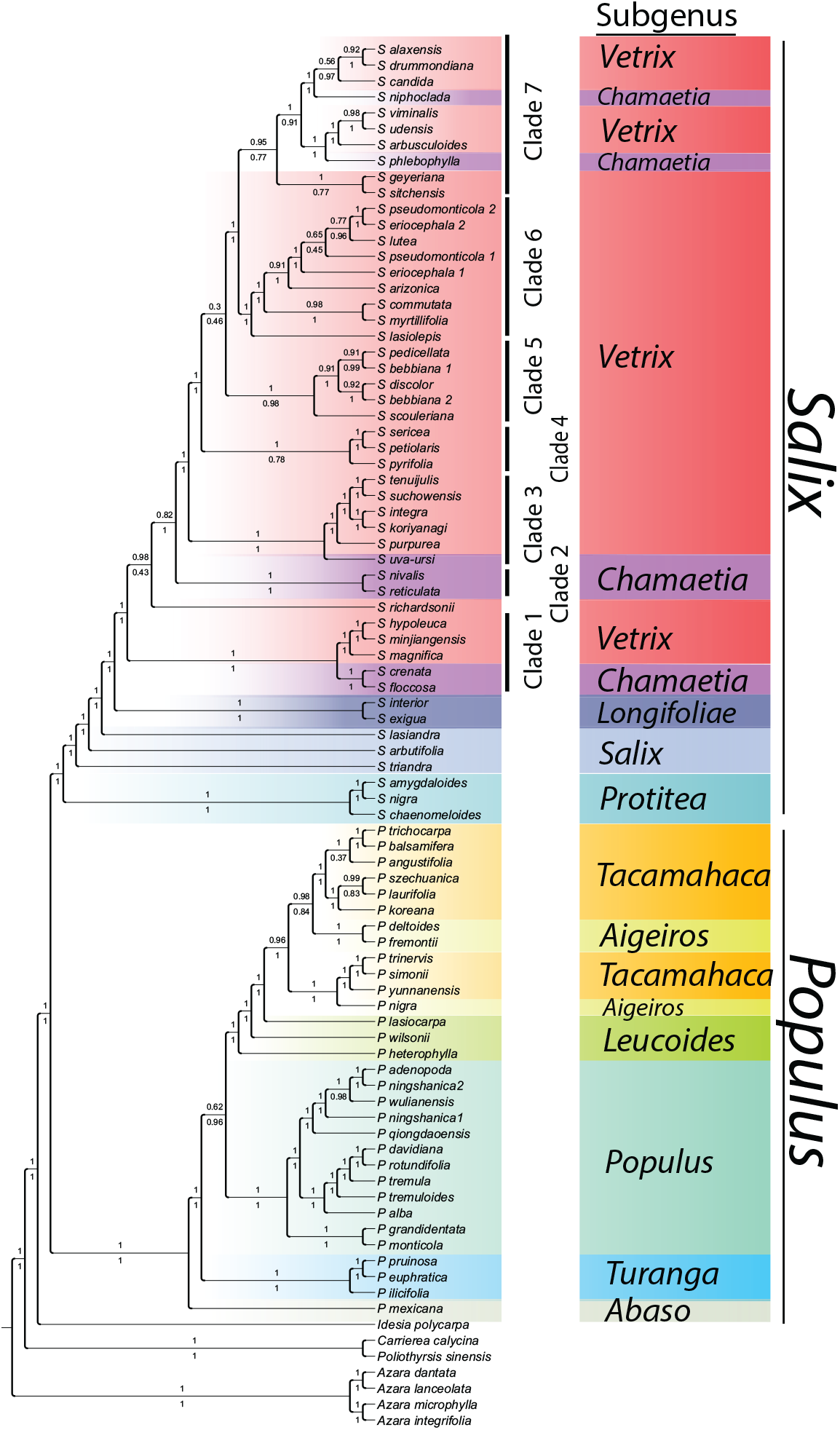
ASTRAL species tree constructed using 787 gene trees. Values above branches indicate posterior probability support and values below branches indicate bootstrap support. Although the most strongly supported relationship among clades is presented, clades with <90% posterior probability or <75% bootstrap support should be viewed with caution. Colors indicate taxon associations with historically identified subgenera.

Most of our samples identified as the same species clustered together in both the ASTRAL tree of individuals and the concatenated tree (Fig. S1). Both the ASTRAL tree and the concatenated tree exhibited high support values for many nodes (especially the concatenated tree). In general, the ASTRAL tree supports the taxonomy of clades that diversified near the root of each genus and provides less support for recently differentiated clades (Fig. S2). The concatenated tree and ASTRAL tree differed in key topological components (Fig. S1) including the placement of *Populus* subgenus *Turanga*. Morphological analyses (Eckenwalder 1996) and a larger set of genes previously analyzed from many of the same *Populus* individuals using only first and second codon positions (Wang et al. 2020), however, generally support a similar topology as our ASTRAL tree (see Discussion). Within *Salix*, the concatenated and ASTRAL trees concurred for the relative placement of subgenera *Protitea, Salix*, and *Longifoliae*, but differed in the placement of groups within *Vetrix* and *Chamaetia*. Within subgenus *Salix* the concatenated and species trees differed in the relative placement of *S. lasiandra, S. triandra*, and *S. arbutifolia*. Finally, there was generally consistent support between the concatenated and species tree for the monophyly of clades within the *Salix+Chamaetia* group with the exceptions of the placement of *S. scouleriana, S. lasiolepis, S. richardsonii, S. geyeriana*, and *S. sitchensis*. Importantly, the ASTRAL tree generally exhibited less bootstrap and SH-alRT support for nodes than the concatenated tree, perhaps indicating that the confidence placed on relationships in the concatenated tree is overestimated. Clades in *Populus* were generally more strongly supported by consistency among gene trees than clades in *Salix* (Figs. 1, S3); this lack of support was especially apparent in the backbone of the *Chamaetia+Vertix* clade. Because conflict among gene trees likely results from either ancient or ongoing gene flow among taxonomic groups (hybridization) or incomplete lineage sorting (ILS), we tested for patterns of historical gene flow and rapid diversification in our tree.

Levels of biased intraspecific gene flow indicative of hybridization were approximately the same in *Populus* and *Salix* (52% of Dtree and 60% of D_min_ were significant after Benjamini-Hochberg adjustment in *Salix* vs. 64% of Dtree and 58% of D_min_ in *Populus* (Tables S4, S5, S6, S7). Because D and f_4_ statistics across a clade are phylogenetically correlated, we used the heuristic f_branch_ analysis based on the ASTRAL species tree to assess the history and timing of hybridization during the diversification of *Populus* and *Salix* (Fig. 2, Table S8, S9). The number of f_branch_ statistics that indicated >5% gene flow across species due to hybridization were similar in *Salix* (6.6% of f_branch_ values above 5%) and *Populus* (7.0% of f_branch_ values above 5%) but were more commonly associated with deep internal branches of *Salix* than *Populus* (Tables S8, S9). Up to three ancient hybridization events appear to have influenced interspecific gene flow in *Salix*. First, the f_branch_ metrics indicated evidence for gene flow between the ancestors of both *S. triandra* and *S. arbutifolia* and ancestors of the lineage leading to subgenera *Longifoliae, Vetrix* and *Chamaetia* (f_branch_ = 0.088 & 0.113; arrow 1 in Fig. 2; Table S8). Second, ancient geneflow (f_branch_ > 0.05%) occurred between the ancestors of species in clade 1 with ancestors of clade 2-7 (f_branch_ = 0.126 to 0.164; arrow 2 in Fig. 2; Table S8). Finally, the f_branch_ analysis indicates ancient hybridization among ancestors of clades 6 and 7 (f_branch_ = 0.078 to 0.186; arrow 3 in Fig. 2; Table S8), as well as significant hybridization between ancestors of clades 6+7 and clades 1-5 (f_branch_ = 0.077 to 0.218; arrow 3 in Fig. 2; Table S8). The lack of evidence of hybridization in *S. geyriana* and strong evidence for hybridization between *S. sitchensis* and an ancestor of clades1-7 suggests that either *S. sitchensis* or *S. geyriana* may be misplaced on the phylogeny and the multiple signals of hybridization result from a single ancient event. In *Populus* the f_branch_ analysis based on the ASTRAL species tree indicated widespread hybridization among ancestors or extant members of the two major clades comprising the *Populus* subgenus (f_branch_ = 0.120 to 0.259; arrows labelled 4 in Fig. 2; Table S9) as well as signals of ancient or ongoing hybridization within the *Tacamahaca/Aigeiros* clade and the *Leucoides* grade (arrows labelled 5 in Fig. 2; Table S9).

**Figure 2.**
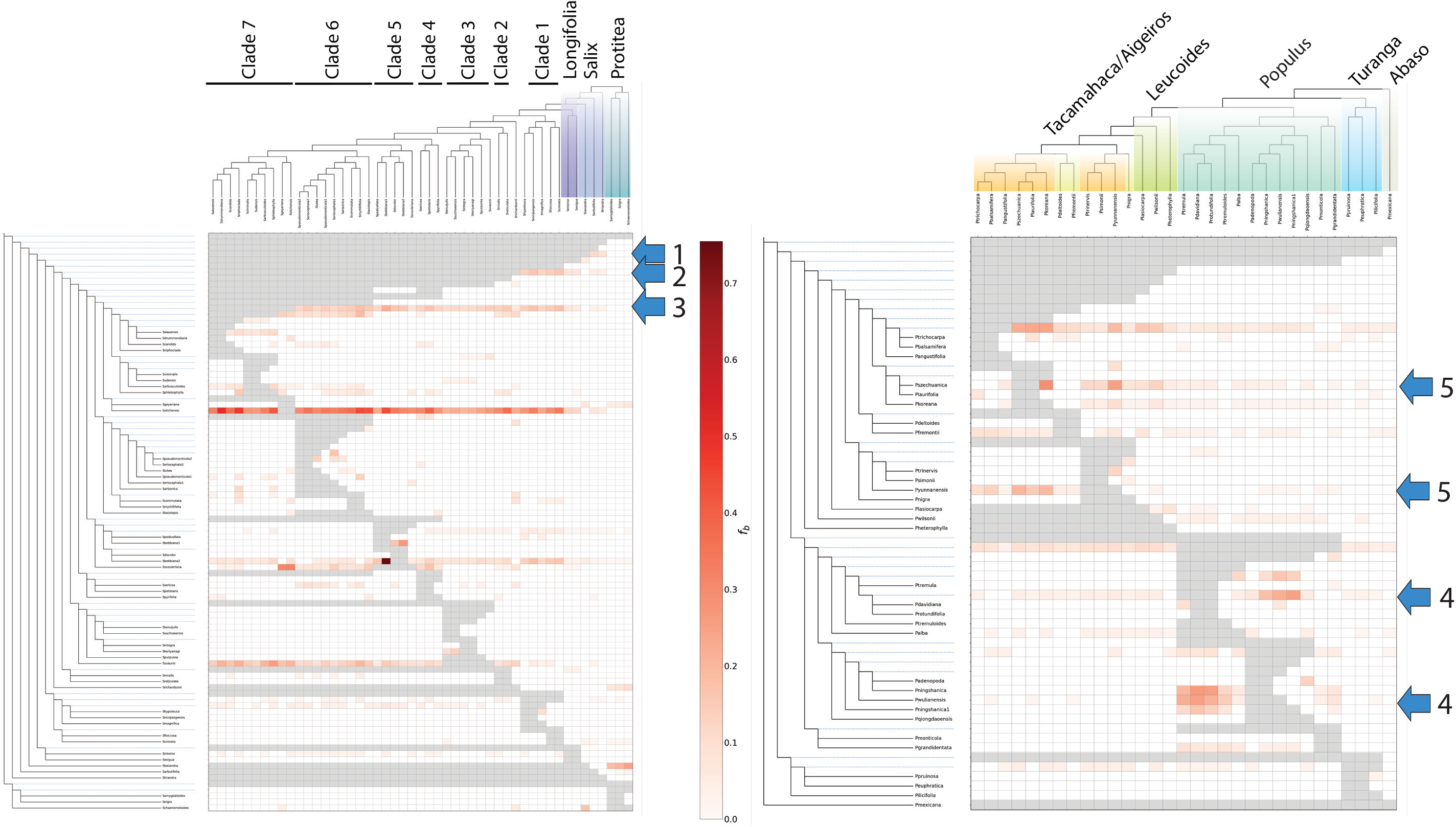
Heuristic f-branch analysis of ongoing and ancient hybridization in Salix and Populus. Arrows indicate gene flow events that are discussed in the text. Color scale in boxes represents the estimated fbranch value, which is indicative of the proportion of the genome that is affected by interspecific gene flow due to hybridization.

The two data sets used to estimate the *BEAST2 dated species trees were both large (>8500 sites) and drawn from genes in the top 7% of our objective criteria. Dating of many nodes was within the overlap of 95% highest posterior density for the two estimates (HPD; Fig. 3; Tables S10, S11). For instance, the divergence between *Populus* and *Salix* was ~35Mya in both data sets (node ‘a’ in Fig. 3), the divergence of *P. mexicana* from the remainder of *Populus* was ~15Mya in both data sets (node ‘d’), and *Salix* subgenera *Vetrix* and *Chamaetia* began to differentiate ~5 mya in both data sets (node ‘c’). Nonetheless, a few estimates of divergence times differed substantially between the data sets with no overlap in 95% HPD. In *Populus* these included the timing of divergence of subgenera *Turanga* (node ‘e’) and *Populus* (node ‘f’) from the remainders of the genus, and in *Salix* these included the timing of divergence of subgenera *Protitea* (node ‘b’) from the remainder of the genus. This inconsistency reflects variance among the gene histories within *Populus* and *Salix* and identify regions of the tree where interpreting the ages of divergence require elevated caution. We also note here that our HPD estimates were constrained by our selection of the standard deviation of the a priori distributions of calibration dates to σ=1. When σ was set to 3, the uncertainty for node dates increased dramatically (Fig. S4; Tables S10, S12).

**Figure 3.**
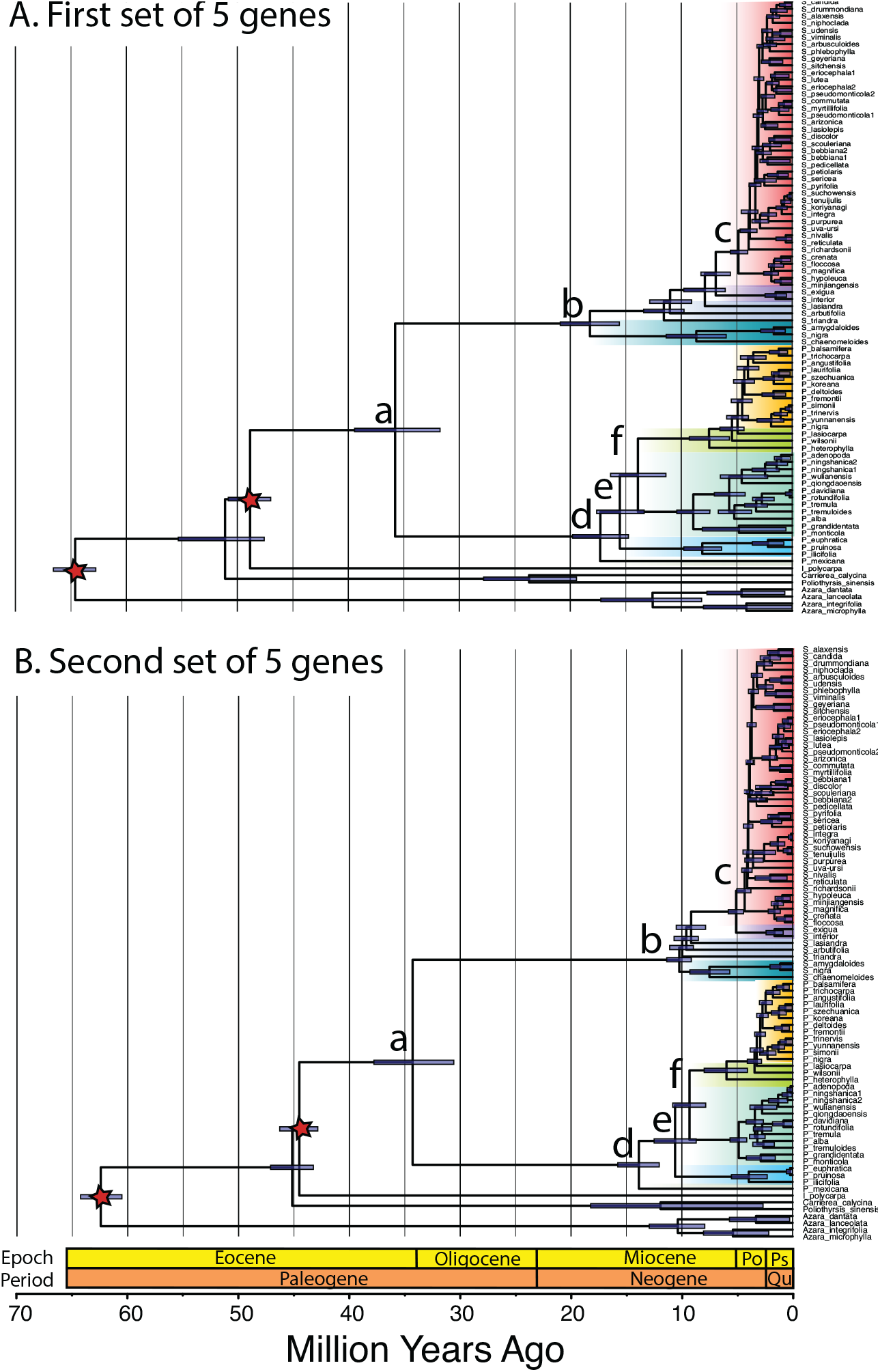
Comparison of dated trees estimated from two different sets of 5 genes using by *BEAST2. A) first set of 5 genes, which was the best set according to criteria described in the methods. B) second set of 5 genes, which was the second-best set. Stars indicate calibration nodes. Node bars represent 95% highest posterior density in node height. Subgenera are colored as in Fig. 1. Letters at nodes are discussed in the text.

Despite slightly different estimates of node ages in our two estimates, both the BAMM and ClaDS analyses indicated strong support for similar patterns of shifts in the diversification rates within both of our dated trees (Fig. 4; Table S13). In both dated trees diversification rate increased near the *Populus-Salix* split and a second increase occurred near the origination of the *Vetrix-Chamaetia* clade in Salix (Fig. 4). For the first dated tree, the BAMM analysis identified two branches with high marginal shift probabilities (Fig. 4A), and these shifts were largely supported in the ClaDS analysis (Fig. 4C). The BAMM analysis also identified substantial support for an increase in diversification rate near the *Populus-Salix* split for the second dated tree, with the exception that the model with most support indicated that *Populus* subgenus *Abaso* (with only *P. mexicana*) retained the same diversification rate at the outgroups (Fig. 4B). It is notable that the marginal odds ratio supporting the shift in diversification at the branch that included all of *Populus* was nearly as high (0.33-0.34) as the branch that did not include *P. mexicana* (0.48-0.50), indicating that the shift was likely near the base of the *Populus + Salix* clade, but may not have included *Populus* subgenus *Abaso*. The ClaDS analysis also supported a rate shift in this general region of the phylogeny. The BAMM analysis also identified in the second dated tree substantial support for an increase in diversification rate in clade that includes the subgenus *Longifoliae* along with the *Vetrix-Chamaetia* clade in *Salix* (Fig. 4B). Again, this pattern was supported in the ClaDS analysis (Fig. 4D).

**Figure 4.**
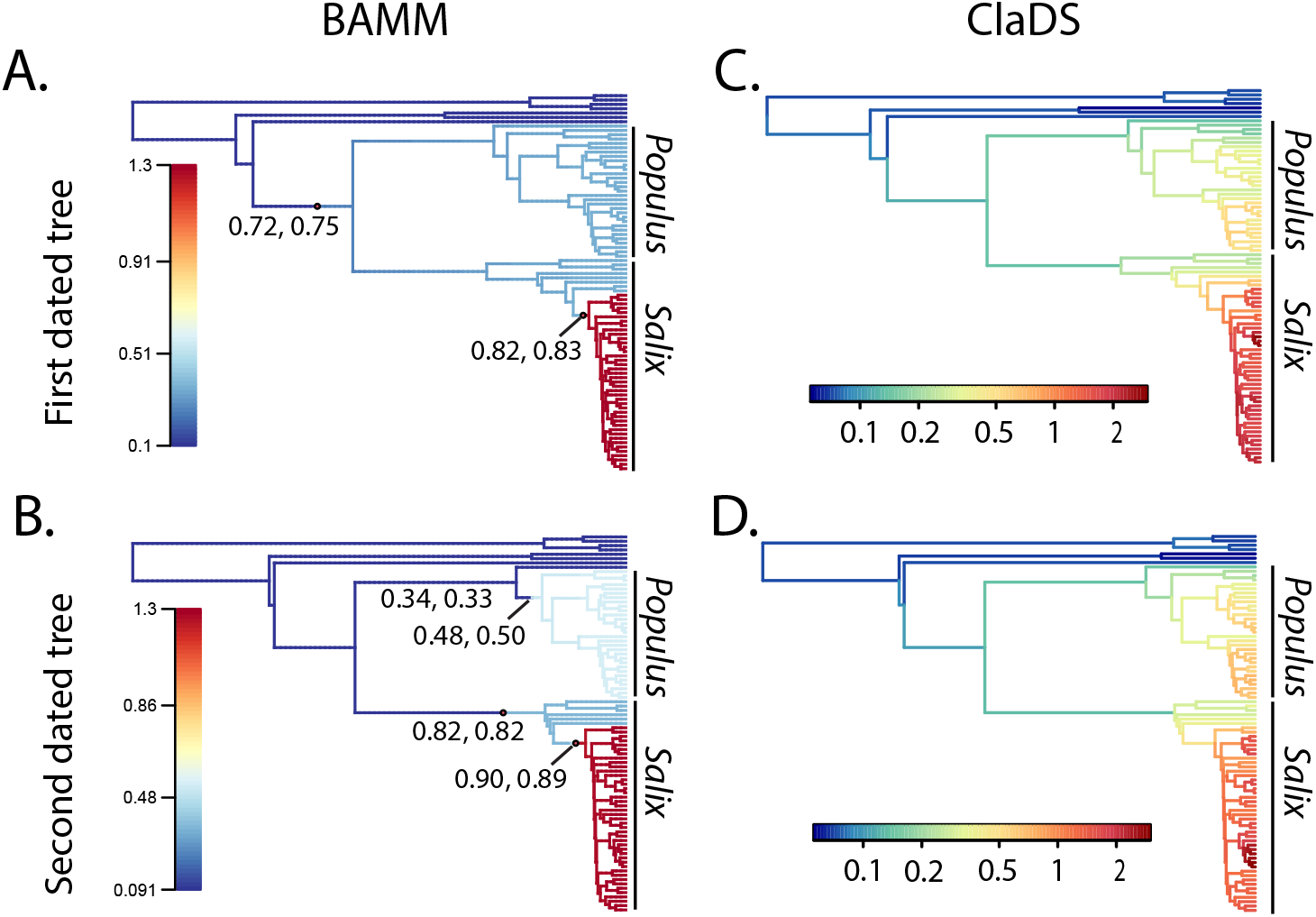
Diversification rate changes across the Salicaceae. Panels A and B show the best estimates of the rate shift set estimated using BAMM for the first and second dated trees, respectively. 60% and 61% (replicate 1 & 2) of samples in the posterior were assigned the the shift configuration shown in A, whereas 40% and 42% (replicate 1 & 2) of samples in the posterior were assigned the the shift configuration shown in B. The 95% credible shift sets for both trees and replicates are presented in Figures S5 and S6. Colors in A and B represent averaged lineage diversification rates according to the scales associated with each panel. Numbers at nodes represent the marginal odds ratios for a shift along branches associated with rate shifts; estimates from the two replicates are presented separated by a comma. Panels C and D show the ClaDS branch-specific rate estimates for the first and second dated trees, respectively. The color scale represents estimate branch specific speciation rates.

We noted that these transitions in diversification rates were associated with shifts in pollination mode (insect in outgroups and *Salix*; wind in *Populus*), seed dispersal mode (animals in outgroups, wind in *Populus* and *Salix*), and growth form (trees in outgroup, *Populus*, and *Salix* subgenus *Protitea;* shrubs and dwarf shrubs in remaining *Salix* subgenera), and a shift from XY to ZW sex determination systems in *Salix* (Sanderson et al. 2021; Gulyaev et al. 2022; Hu et al. 2022). However, there are too few phylogenetically independent trait transitions on our Salicaceae phylogeny to provide a statistically robust test of the associations between traits and diversification rates. STRAPP analyses resulted in no statistically significant associations (P>0.30 for all tests).

## Discussion

The scale of hybridization on *Populus* and *Salix* is remarkable in its taxonomic, genomic, and chronological extent. Signals of ancient hybridization and introgression across diverged lineages have persisted in descendant lineages, are particularly evident in *Salix* and may better account for gene tree discordance than contemporary hybridization. Our f_branch_ analysis estimated that approximately 10-20% of the *Salix* genomes in the *Vetrix-Chamaetia* clade were affected by the persistence of genes from ancient hybridization events, which based on our dated trees, may have begun as early as 5 Mya. This pattern is consistent with the low levels of chloroplast genomic diversity across the *Vetrix-Chamaetia* subgenera which was hypothesized to have been partially influenced by high levels of hybridization and introgression (Wagner et al. 2021b). Earlier hybridization events affecting genomic variation in subgenera *Salix* and *Longifolae* likely occurred even earlier, perhaps nearly 10Mya. These hybridization events have likely contributed to the difficulties in reconstructing relationships within *Salix* in the present and previous studies (Leskinen et al. 1999; Barkalov et al. 2014; Percy et al. 2014; Lauron-Moreau et al. 2015; Wu et al. 2015; Liu et al. 2016). In *Populus*, signals of hybridization between subgenus *Turanga* and several members of *Aigieros-Tacamahaca* likely contributed to the low support for the position of subgenus *Turanga* clade and may underlie incongruence between the ASTRAL and concatenated trees (Fig. S1). Nonetheless, caution is required when interpreting the f_branch_ analysis because it is dependent on the correct topology of the framework phylogeny. An example of this impact is evident in the high level of hybridization indicated between *S. sitchensis* and most other species in the *Vetrix-Chamaetia* clade, yet the lack of hybridization between the putative close relative *S. geyeriana* and these same taxa. This pattern was unlikely to have resulted from recent hybridization between *S. sitchensis* and each one of these species but more likely resulted from the misplacement of *S. sitchensis* and/or *S. geyeriana* in the phylogeny.

Previous studies have reported signals of ancient hybridization in both genera. Two ancient chloroplast capture events have been previously identified in *Populus*, one in which an ancestor of *P. heterophylla* captured the chloroplast of *P. mexicana* ancestors (Liu et al. 2017; Wang et al. 2020) and a second in which an ancestor of *P. nigra* captured the chloroplast of *P. alba* ancestors (Smith et al. 1990). Our data, however, found no evidence of ancient hybridization among these lineages in the nuclear genomes. This lack of evidence may result from the limited numbers of genes that we sampled, or these ancient hybridization events may have only influenced the chloroplast genomes (Tsitrone et al. 2003). Chloroplast capture reflecting hybridization of ancient lineages also has been reported in *Salix* (Brunsfeld et al. 1992; Hardig et al. 2000; Gulyaev et al. 2022). An ancestral member of subgenus *Longifoliae* captured a chloroplast of a member of subgenus *Protitea* (Gulyaev et al. 2022) and five cases of arguably more recent chloroplast capture within *Salix* section *Longifoliae* were reported by Brunsfeld et al. (1992). Notably, crossing studies indicate that extant members of subgenera *Longifoliae* and *Protitea* are reproductively isolated (Mosseler 1990), so the hybridization generating the former chloroplast capture event likely occurred before reproductive isolation evolved. The deep history of hybridization in *Salix* also is reflected in the evidence for gene flow among ancestral lineages in our study and indicates a long history of concomitant hybridization and speciation in the genus. As others have argued for *Populus* (Cronk et al. 2018), this pattern of ongoing speciation with hybridization may be best represented as a syngameon (Lotsy 1925) and may exhibit emergent properties such as the ability to draw on elevated levels of standing variation for adaptive evolution (Cannon et al. 2020; Cannon 2021). Envisioning *Salix* as a syngameon redefines evolutionary units as larger combinations of hybridizing species and our data and previous results (Hardig et al. 2000; Murphy et al. 2022) suggest that the *Salix* syngameon exhibits a complex web of ongoing hybridization and partial reproductive isolation that has persisted for millions of years.

Although our recovered topology for *Populus* was similar to Wang et al. (2020), it was not identical. A notable difference between our tree and that of Wang et al. (2020) is that our tree placed *P. angustifolia* as sister to the *P. trichocarpa-P. balsamifera* clade, whereas the Wang et al. (2020) placed *P. angustifolia* at the base of a larger clade including multiple North American and Asian species. Importantly, the placement of Wang et al. (2020) indicates that either *P. szechuanica, P. laurifolia*, and *P. koreana* or *P. angustifolia* may have speciated due to vicariance or long-distance dispersal from North America to Asia and the divergence of *P. angustifolia* predates this event. The difference in the placement of *P. angustifolia* is particularly interesting and worthy of further study because it commonly hybridizes with both *P. trichocarpa* and *P. balsamifera* (Brayshaw 1965; Chhatre et al. 2018), which may influence patterns of gene tree coalescence, and the perceived relationships among species. Interestingly, some foliar pathogens of *P. angustifolia* are related to foliar pathogens of Asian members of *Tacamahaca* and absent in North American members (Busby et al. 2012), lending support to the hypothesis of a recent trans-Beringian migration of *P. angustifolia*.

The estimates for divergence times among subgenera within *Populus* and *Salix* presented here must be considered tentative because we did not calibrate internal nodes in each genus. Many fossils of *Populus* and *Salix* from both Asia and North America have been identified (Collinson, M.E. 1992), but it has been difficult to accurately assign them to extant taxonomic groups without robust phylogenies. We chose to rely primarily on the molecular clock to estimate diversification dates within these genera instead of including internal calibrations following Percy et al. (2014) and Zhang et al. (2018). We note that the estimates of diversification dates by Wu et al. (2015) were much earlier than we report here and included an internal *Salix* late-Oligocene (ca. 26Ma) leaf fossil to calibrate of the origination of subgenus *Vetrix* (Wolfe pers. comm. 1991 in Collinson 1992). Also, *Populus* fossils hypothesized to belong to subgenus *Tacamahaca* are described from the Late Oligocene Creede flora of Colorado (Wolfe et al. 1990; Collinson 1992), a date much earlier than our estimate of the diversification of this subgenus. However, our and other recent phyogenies have identified both subgenus *Tacamacaha* and *Vetrix* as polyphyletic (Wagner et al. 2018; Wang et al. 2020), suggesting that the characters used to categorize the fossils should be reexamined. The last thorough review of Salicaceous fossils was published in 1992 (Collinson 1992), and updating of this group in relation to the most up-to-date phylogenies would aid greatly in developing a more accurate estimate of diversification times in the Salicaceae.

In *Salix* we identified a burst of diversification near the origin of the *Vetrix-Chamaetia* clade that likely increased the level of incomplete lineage sorting (ILS) and contributed to the lack of support for inferred relationships within this clade (Roch et al. 2015). We also identified a second increase in the diversification rate near the divergence of *Populus* from *Salix*. The mechanisms driving these increases in diversification rate remain speculative. Although these shifts are accompanied with changes in seed dispersal, pollination vectors, and growth form (Argus 2010; Eckenwalder 2010), and even a shift from an XY to a ZW sex determination system in *Salix* (Gulyaev et al. 2022; Hu et al. 2022), each of these shifts occurs only once or twice on the tree, so too few phylogenetically independent events have occurred for powerful statistical tests of association. Nonetheless, these patterns may be useful for meta-analyses including a larger set of taxonomic groups, or future studies may find more detailed patterns associated with genetic or morphological changes that can shed light on the drivers of diversification in these two important genera.

The continued development of analytical methods for quantifying ancient interspecific gene flow will permit investigations into the prevalence of ancient hybridization, its impacts on adaptation, and its biological and phylogenetic correlates. Ancient hybridization inferred from chloroplast capture has been identified in large numbers of plants groups (Rieseberg et al. 1991). However, chloroplast capture may result from unique properties of the hybridizing species (Tsitrone et al. 2003) and may not be present in all taxonomic groups affected by ancient hybridization. Interspecific gene flow is usually not categorized as either contemporary or ancient, but the relative influences of ancient versus recent introgression events may provide insight into the sources of different classes of genetic variants under selection (e.g. Menon et al. 2021). Recent studies in *Quercus, Pinus*, and *Populus* have documented adaptive introgression among hybridizing groups of multiple species (Chhatre et al. 2018; Leroy et al. 2020; Buck et al. 2022). Along with *Populus* and *Salix*, ancient gene flow has also been identified in *Quercus*, where phylogenetic conflict in chloroplast genomes suggests hybridization during early diversification (Yang et al. 2021) indicating at least two plant families with a combination of both contemporary and ancient interspecific gene flow. Plant families differ in their propensity to hybridize, and there is a strong phylogenetic signal for hybridization (Whitney et al. 2010). Thus, it is likely that the ancestors of contemporary groups with strong propensities for hybridization also hybridized. Understanding patterns of ancient interspecific gene flow in relation to contemporary hybridization may provide further insight into biological factors correlated with hybridization (Mitchell et al. 2019) and factors associated with the development of syngameons.

## Conclusion

Examples of the extended effects of contemporary hybridization in *Salix* and *Populus* are well-known (Hardig et al. 2000; Evans et al. 2008; Lexer et al. 2010). The cumulative effects of persistent hybridization over eons as diversification unfolded, however, may result in different qualities of adaptation and diversification than isolated cases of contemporary hybridization. In *Salix* and *Populus* this history has resulted in a tangle of gene histories within each genus, with some clades having developed monophyly and others that may never be resolved. Hybridization also may have contributed changes in the diversification rate of species in the *Vetrix-Chamaetia* clade of *Salix*. Understanding the characteristics associated with and generated from long-term and persistent interspecific gene flow will elucidate whether these properties confer fundamentally different pattens of adaptation and speciation (Cannon 2021).

## Data Accessibility

All alignments and gene trees used in this research are available at: https://datadryad.org/stash/share/48HL7Z3TAgH1hOQLwWM1EzundQOFxN0fwoRxXMdXi_s All data not already available at NCBI will be uploaded to the NCBI short read archive prior to publication – we are currently finalizing the metadata.

## Acknowledgements

We thank Peter Zhelev and Gancho Slavov for *S. triandra* collections, Andrew Hamstetter for insightful discussions of diversification analyses, and Pascal Title for computational assistance with BAMM. This research was supported by grants from the US National Science Foundation (1542509, 1542599, 1542479, 1542486) and the National Natural Science Foundation of China (31561123001).

