## Supplemental Methods for "Phylogenomics reveals patterns of ancient hybridization and differential diversification contributing to phylogenetic conflict in *Populus* L. and *Salix* L"

**Supplemental methods for "Phylogenomics reveals patterns ancient hybridization and differential diversification that contribute to phylogenetic conflict in *Populus* L. and *Salix* L."**

Sanderson et al. (2020) details the development of the sequence capture kit targeting 972 genes (average exon length 1098 ± 489 bp) using 12,951 probes used in this study. A subset of these genes were involved in the Salicoid whole genome duplication, and so our reference for assembling the target capture reads included an additional 247 paralogs that also may be captured (1219 genes total). This sequence capture kit was used to genotype 14 *Populus* samples (7 species) and 83 *Salix* samples (45 species) for this study. DNA was extracted from silica gel dried leaf tissues using CTAB or Qiagen DNeasy Plant kits. Libraries were constructed and captured using methods and probes (Daicel Arbor Biosciences Ref #170424-30 "Salicaceae") outlined in Sanderson et al. (2020). Primers and low quality bases were trimmed from read data using Trimmomatic v 0.36 (Bolger et al. 2014). Whole genome sequences from 69 additional samples were added to the data set; some of these were phylogenetically analyzed here for the first time and others analyzed in former studies (Table S1).

Sequence data from 166 samples were assembled into putatively homologous gene sequences using the HybPiper pipeline (Johnson et al. 2016). Paralogous sequences may align to the same reference gene, with the potential to generate trees that are inconsistent with the species tree or have excessively long branches. To address these concerns, 413 sequence capture locus alignments that included potential paralogs were identified using Hybpiper, and both copies were removed from the final data set. These paralogs were defined as genes/loci with multiple contigs and comprising 80% of the length of the target region (Johnson et al. 2016). Amino acid sequences recovered from HybPiper were aligned using UPP (Nguyen et al. 2015b) and converted back to codon-aligned nucleotide sequences using PAL2NAL v 14 (Suyama et al. 2006). For each gene sequence, all sites with >25% gaps (-gt 0.75) and without at least 90% of residues that overlap of at least 90% with the rest of the sequences (-resoverlap 0.90 -seqoverlap 0.90) were removed using trimal v1.4.rev22 (Capella-Gutierrez et al. 2009). TreeShrink 1.3.4 was used to detect and filter out sequences with excessively long branches in each gene tree (Mai et al. 2018). 787 alignments remained after filtering and removing gene alignments likely containing paralogous sequences and were used for all downstream phylogenetic analyses (Table S2).

For the ASTRAL approach, trees for each of the 787 genes were estimated using IQTREE 2.0.3 (Nguyen et al. 2015a) with the best substitution models identified for each tree independently (-MFP; Kalyaanamoorthy et al. 2017) and 1000 multilocus bootstrap replicates per tree (Hoang et al. 2018). For each gene tree, branches were collapsed when bootstrap support was <33% using sumtrees in the DendroPy package (Sukumaran et al. 2010) prior to ASTRAL analysis. The ASTRAL tree of all individuals and the species tree (Rabiee et al. 2019) were inferred using ASTRAL-MP v 5.12.2 (Yin et al. 2019). Trees were plotted using TreeGraph 2 (Stover et al. 2010) or FigTree 1.4.4 (http://tree.bio.ed.ac.uk/software/figtree/). Gene tree concordance with the best ASTRAL tree was estimated using phyparts (Smith et al. 2015) and plotted using phypartspiecharts (github.com/mossmatters/phyloscripts).

The dated ultrametric species trees were calculated using *BEAST2 (Heled et al. 2010). Our best estimate was based on information from 5 genes using all individuals (9,264 sites: SapurV1A.0003s0350, SapurV1A.0045s0240, SapurV1A.0050s0650, SapurV1A.0139s0330, SapurV1A.0260s0050). The 5 genes were selected in a two-step process that first used SortaDate

(Smith et al. 2018) to identify 40 gene trees with the highest consistency with the species tree topology, minimal root-to-tip variance (most clock-like), and maximum tree length (information content; Smith et al. 2018). In a second step, 5 genes were chosen with phylogenies (estimated using IQTREE) that were visually congruent in gene tree topology with basal nodes of *Populus* and *Salix* in the ASTRAL species tree. To assess the variance of the node age estimates, a second set of 5 genes was selected from the 40 gene trees which scored somewhat lower on the above metrics (8,568 sites: SapurV1A.0211s0160, SapurV1A.0789s0070, SapurV1A.0857s0020, SapurV1A.0900s0040, SapurV1A.1178s0060). For both *BEAST2 analyses the substitution models were linked for all 5 genes, but clock models and trees were unlinked, and the topology was constrained at many nodes to mirror the ASTRAL species tree (constrained nodes identified in Fig. 3). Node dates were modelled using a random local clock under the lineage Birth-Death process using a GTR+Gamma+6 Invariant sites substitution model while estimating base frequencies. For both sets of genes, we ran 5 separate replicate searches with different starting seeds, all with chain lengths of 500M, and sampling every 25000 generations. After trace inspection using Tracer v1.7.2 (Rambaut et al. 2018), 15%-60% of the reads were removed for burn-in depending of the replicate. For both gene sets, results were similar across replicates and were combined for final analyses, which resulted in ESS>200 for most statistics (Tables S10, S11, S12). Finally, to address the impact of the narrow standard deviations around the fossil calibration dates, we re-ran the *BEAST2 analysis for the first set of genes with the same parameters except with an increase in the standard deviation around the calibration dates from 1.0 to 3.0. Results of these analyses are shown in figure S4.

We calculated ABBA-BABA, $f_4$, and $f_{branch}$ statistics from vcf files using Dsuite (Malinsky et al. 2021). We called two separate vcf files using bwa v0.7.17 (Li et al. 2009). One vcf included all of *Populus* samples and *S. nigra* as an outgroup, with *Populus trichocarpa* v 4.1 as a reference (Tuskan et al. 2006). The second vcf included all *Salix* samples and *P. monticola* as an outgroup, with S. purpurea v1.0 genome as a reference (Zhou et al. 2018). The HalpoCaller module from GATK v4.2.6.1 (Van der Auwera et al. 2020)was used to call variants. Filtering was performed using the following criteria: MQ < 45, QD < 5, FS > 60, MQRankSum < -12.5, NS ReadPosRankSum < -8. Final genotypes were filtered for 8<DP<105 for the data generated from whole genome sequencing and 10<DP<1065 for data generated form sequence capture.
