## Supplemental Figures for "Phylogenomics reveals patterns of ancient hybridization and differential diversification contributing to phylogenetic conflict in *Populus* L. and *Salix* L"

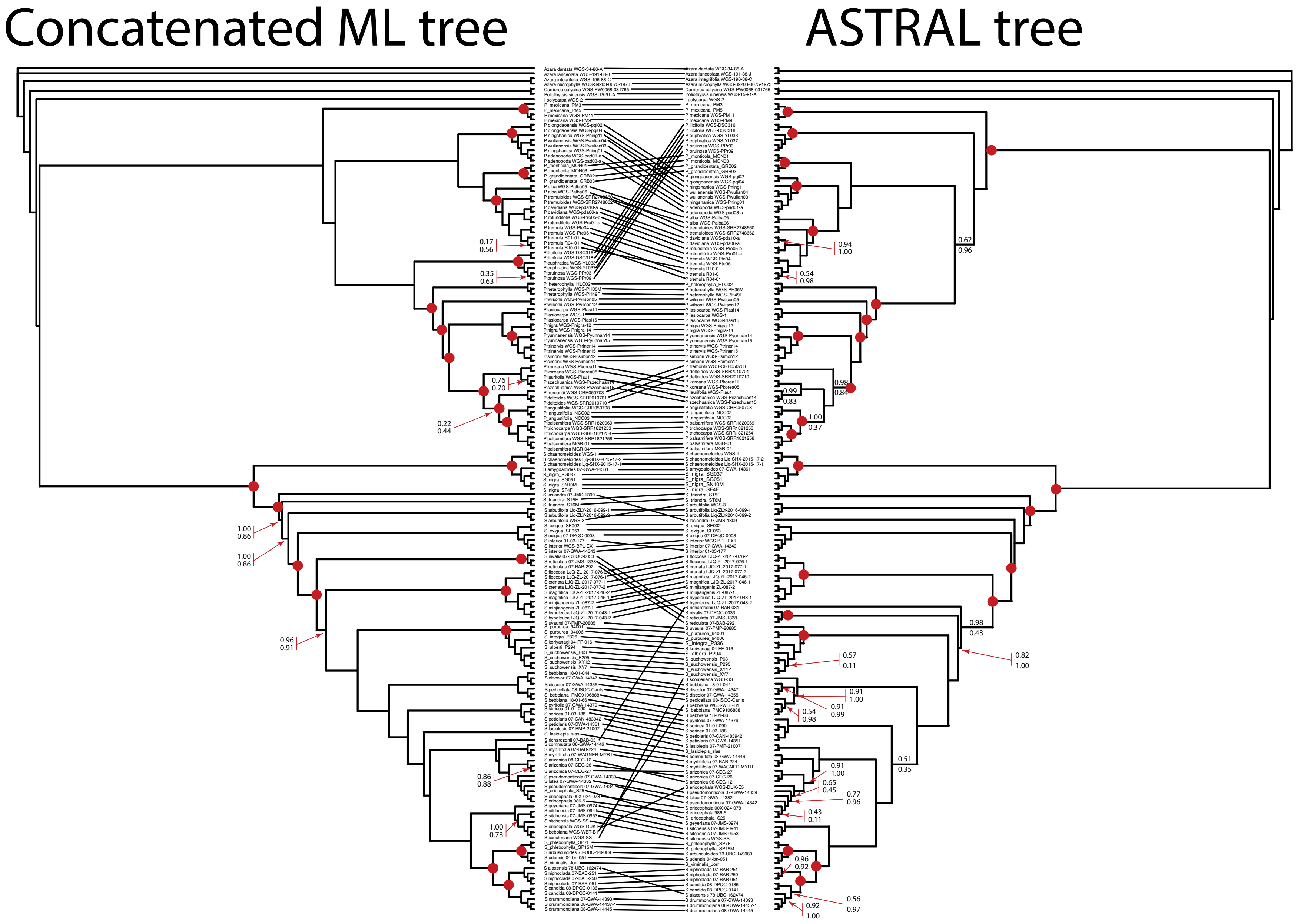


Figure S1. Comparison of the concatenated tree and the Astral tree of of all individuals. Nodes marked with red dots are shared between the trees. Tree support for the concatenated tree is provided as percentage bootstrap/SH-alRT test. Tree support for the super tree is provided as local posterior probability/percentage bootstrap. Unlabeled nodes had >0.95/>0.95 for the relevant metrics.

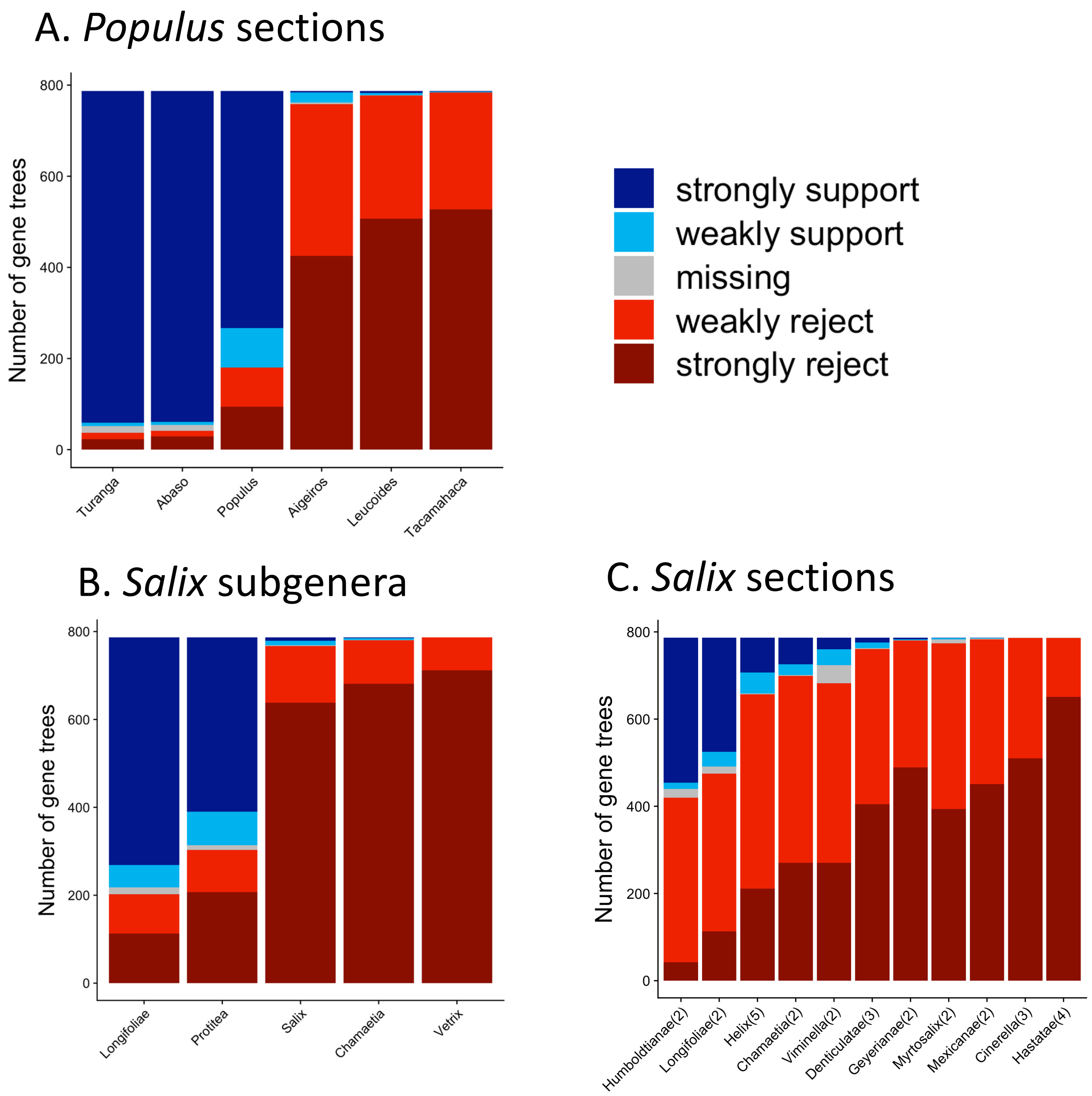


Figure S2. Numbers of gene trees (out of 787) consistent with monophyly of A) *Populus* sections, B) *Salix* subgenera, and C) *Salix* sections. Strongly support indicates monophyletic groups are supported with bootstrap support >75%; weakly support indicates monophyletic groups with <75% bootstrap support. Weakly rejected indicates groups that are not in the tree but are compatible if branches with <75% bootstrap support are collapsed. Strongly rejected indicates groups with no support for monophyly.

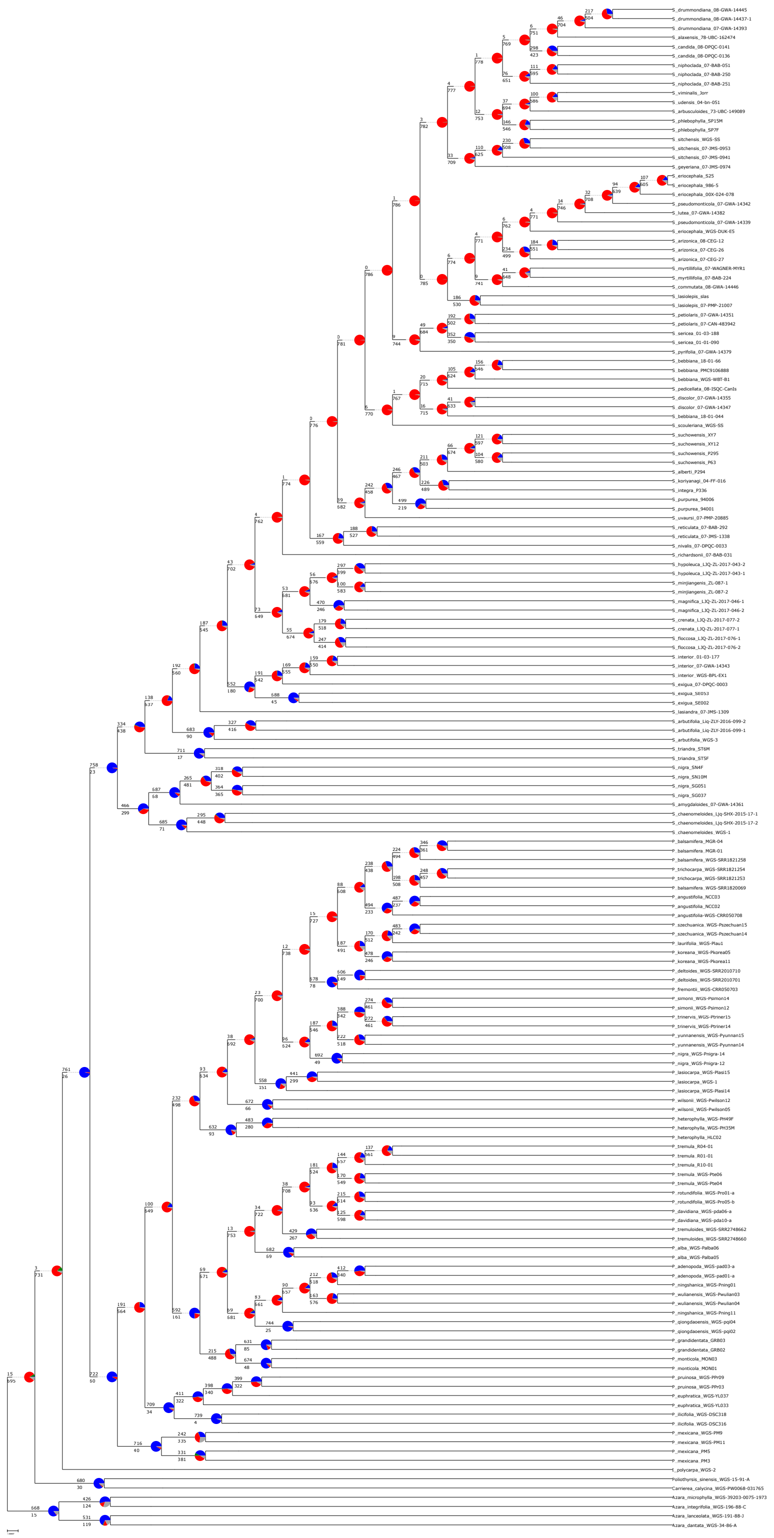

Figure S3. Astral super tree including all individuals constructed using 787 gene trees. Pie charts indicate the number of gene trees consistent with the node (blue), and inconsistent with the node (red). Green indicates a node with an alternative clade that was supported by multiple gene trees that was not present in the best ASTRAL tree shown here.

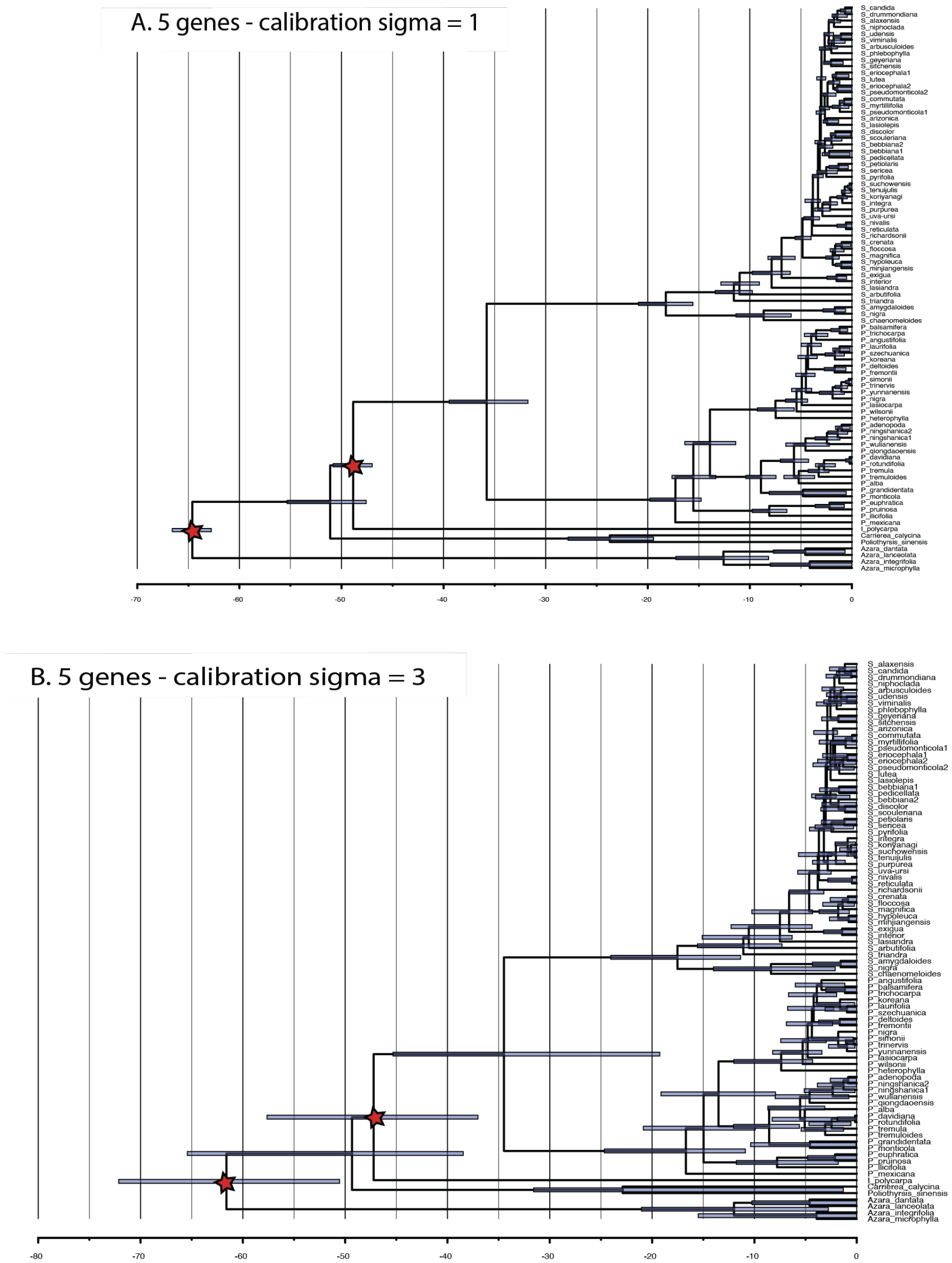


Figure S4. Comparison of ultrametric trees when *a priori* distributions of calibration dates were drawn from a distribution with sigma= 1 (A) and sigma=3 (B).

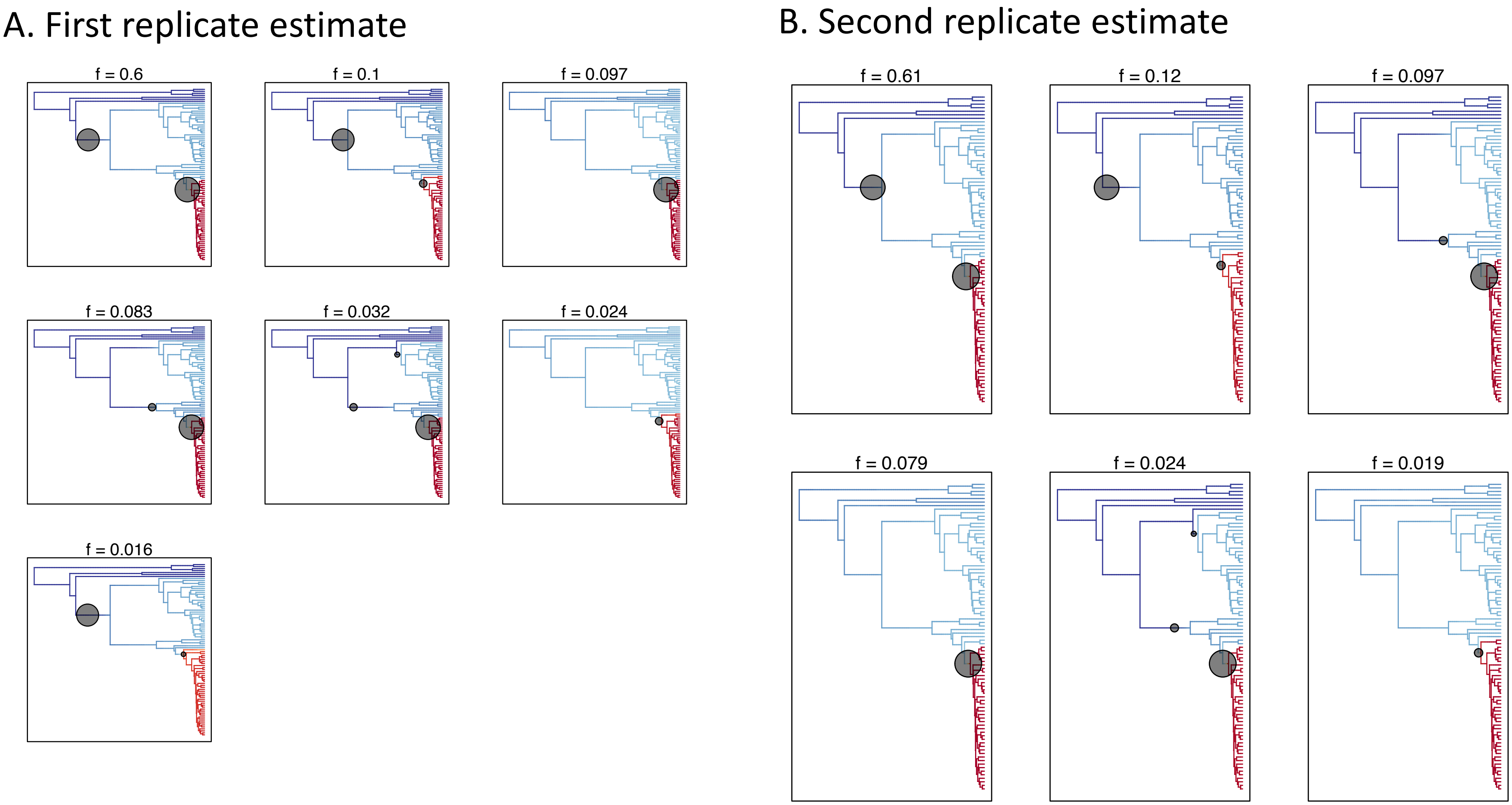


Figure S5. The 95% credible shift sets for changes in diversification rate along the first dated species tree using the dated tree without internal generic calibrations based on the first five selected genes (Fig. 3A). Panel A shows the results from the first replicate run and panel B shows the results from the second replicate run. "f=" indicates the proportion of samples in the posterior were assigned to the shift configuration. Colors indicate averaged lineage diversification rates.

### A. First replicate estimate

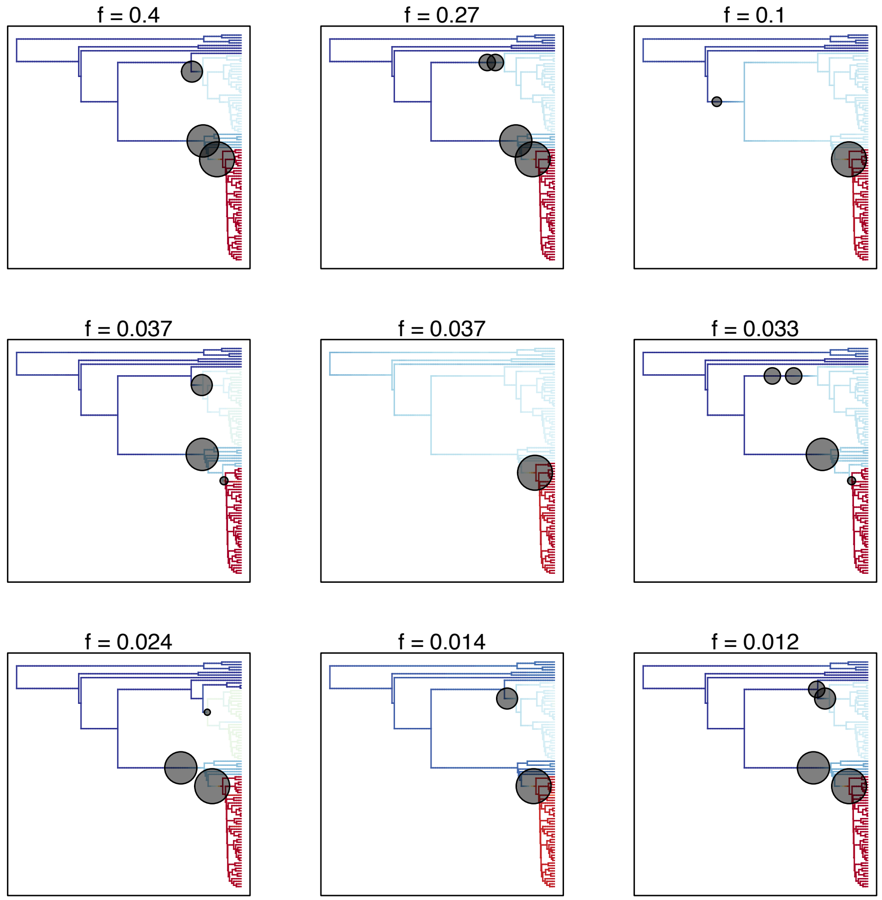


### B. Second replicate estimate

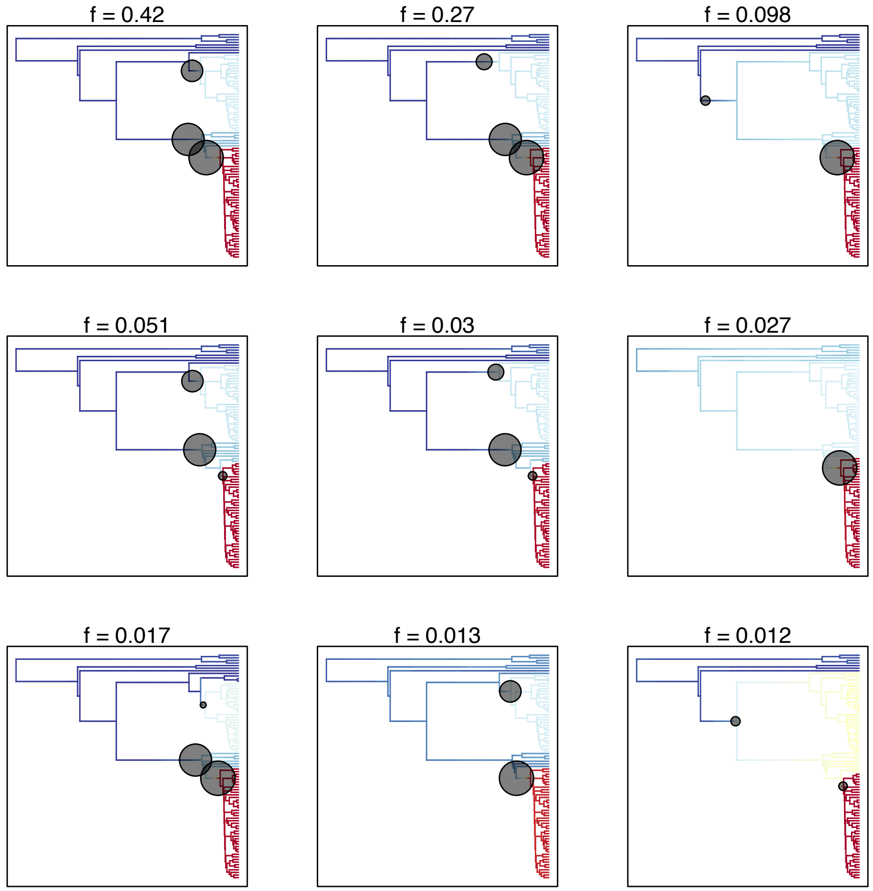


Figure S6. The 95% credible shift sets for changes in diversification rate along the second dated species tree using the ultrametric tree without internal generic calibrations. Panel A shows the results from the first replicate run and panel B shows the results from the second replicate run. “f=“ indicates the proportion of samples in the posterior were assigned to the shift configuration. Colors indicate averaged lineage diversification rates.

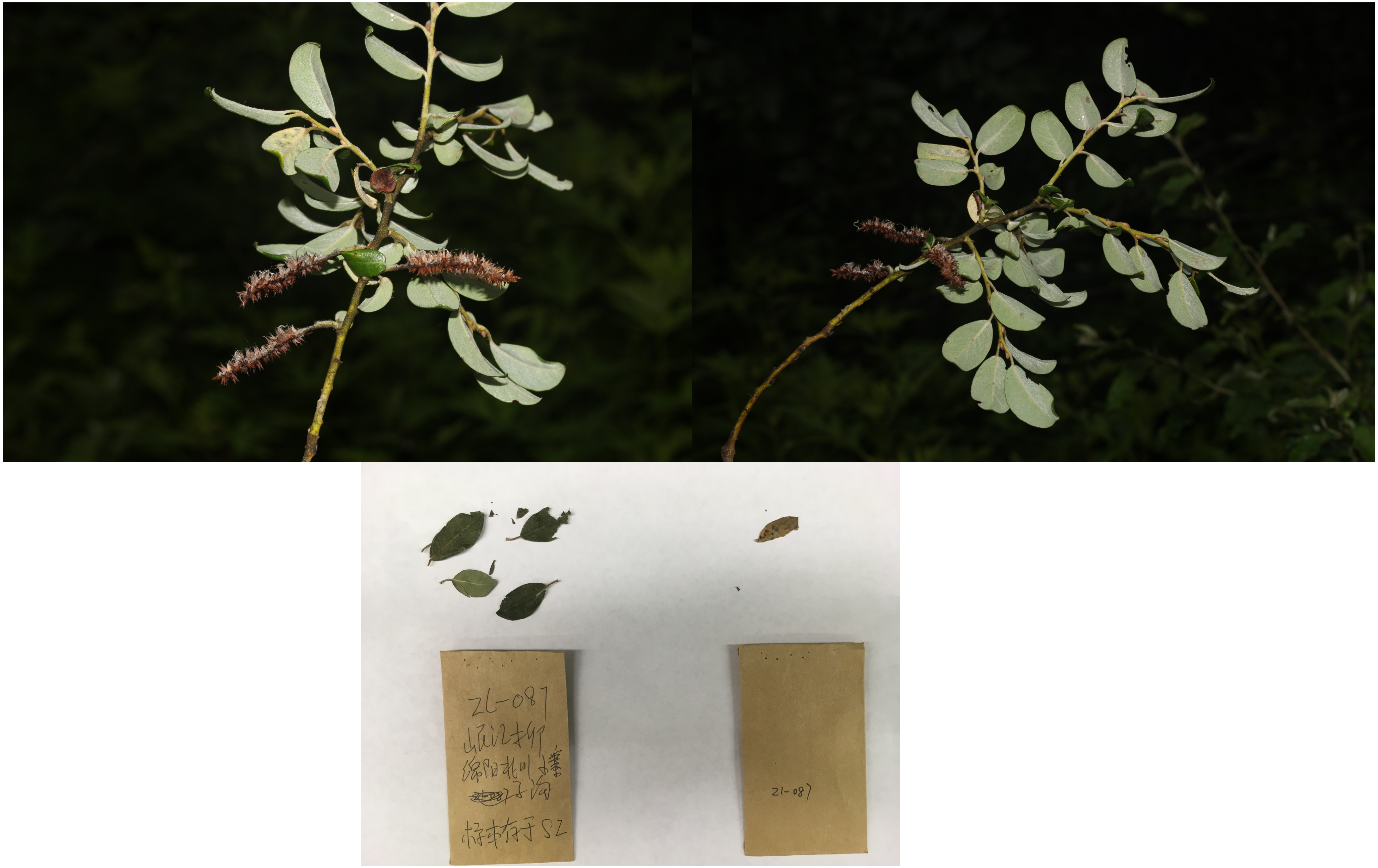


Figure S7. Photographic vouchers for specimens. a. *Salix mingjiangensis*

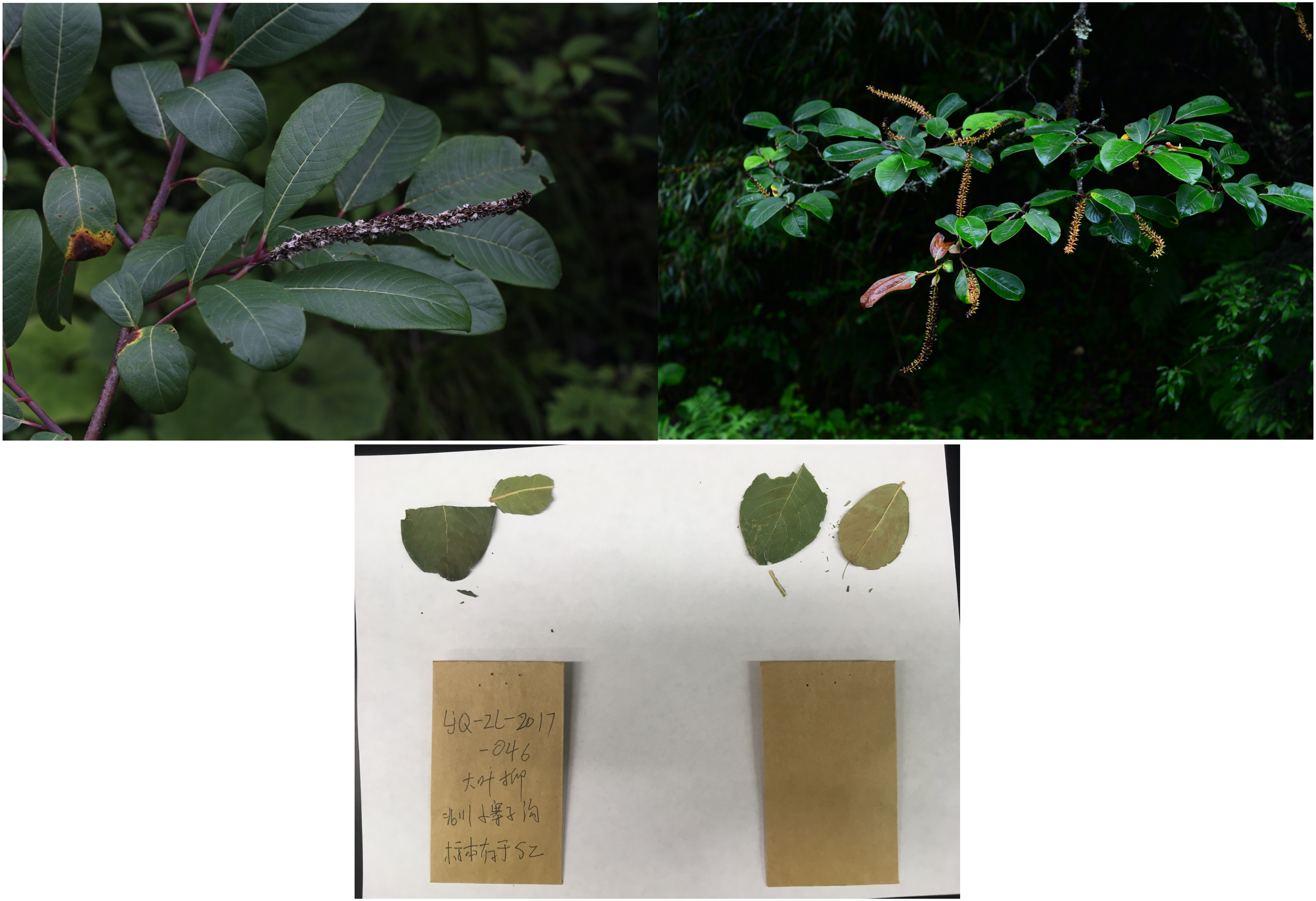


Figure S7. Photographic vouchers for specimens. b. *Salix magnifica*

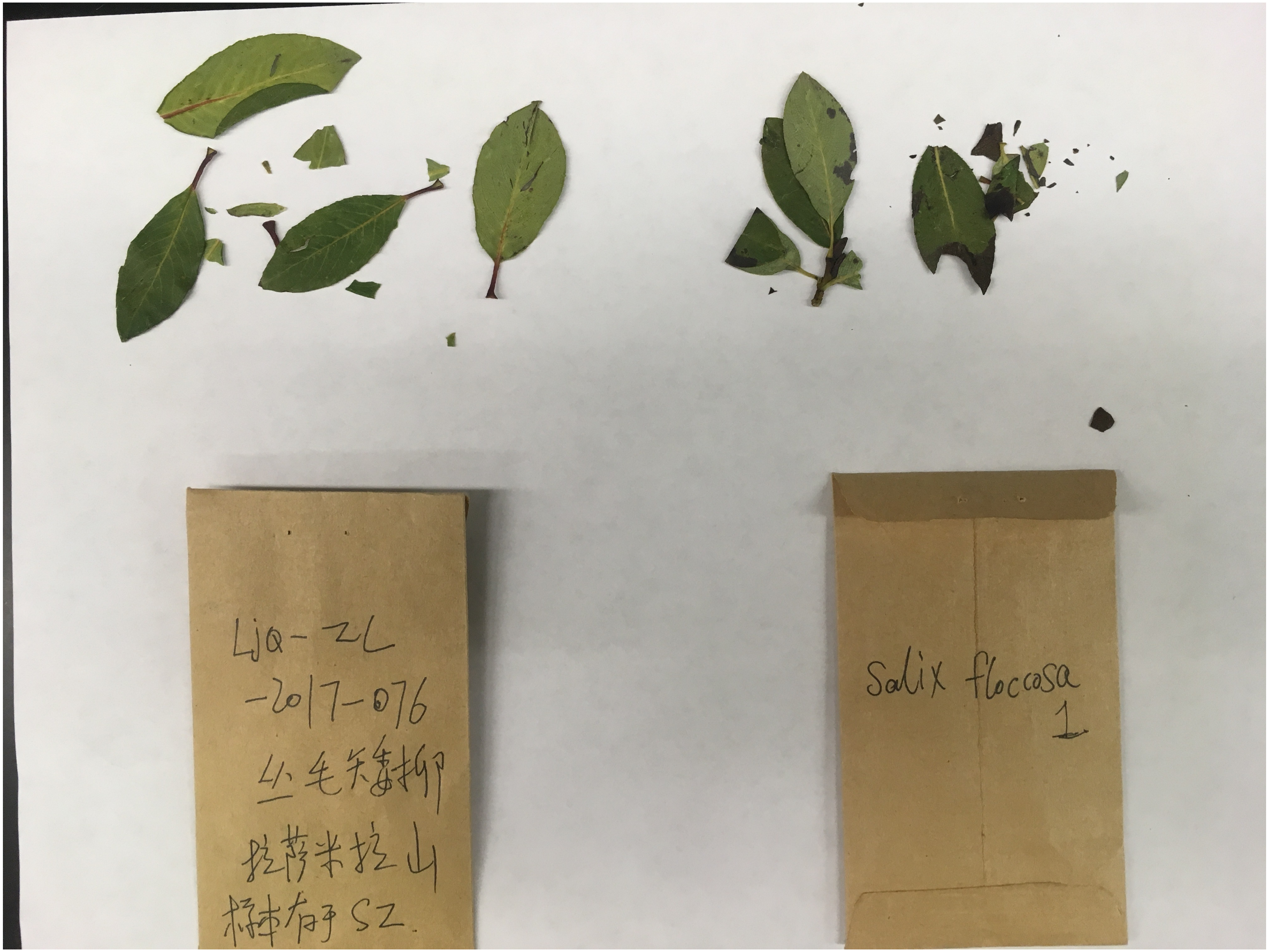


Figure S7. Photographic vouchers for specimens. c. *Salix flaccosa*

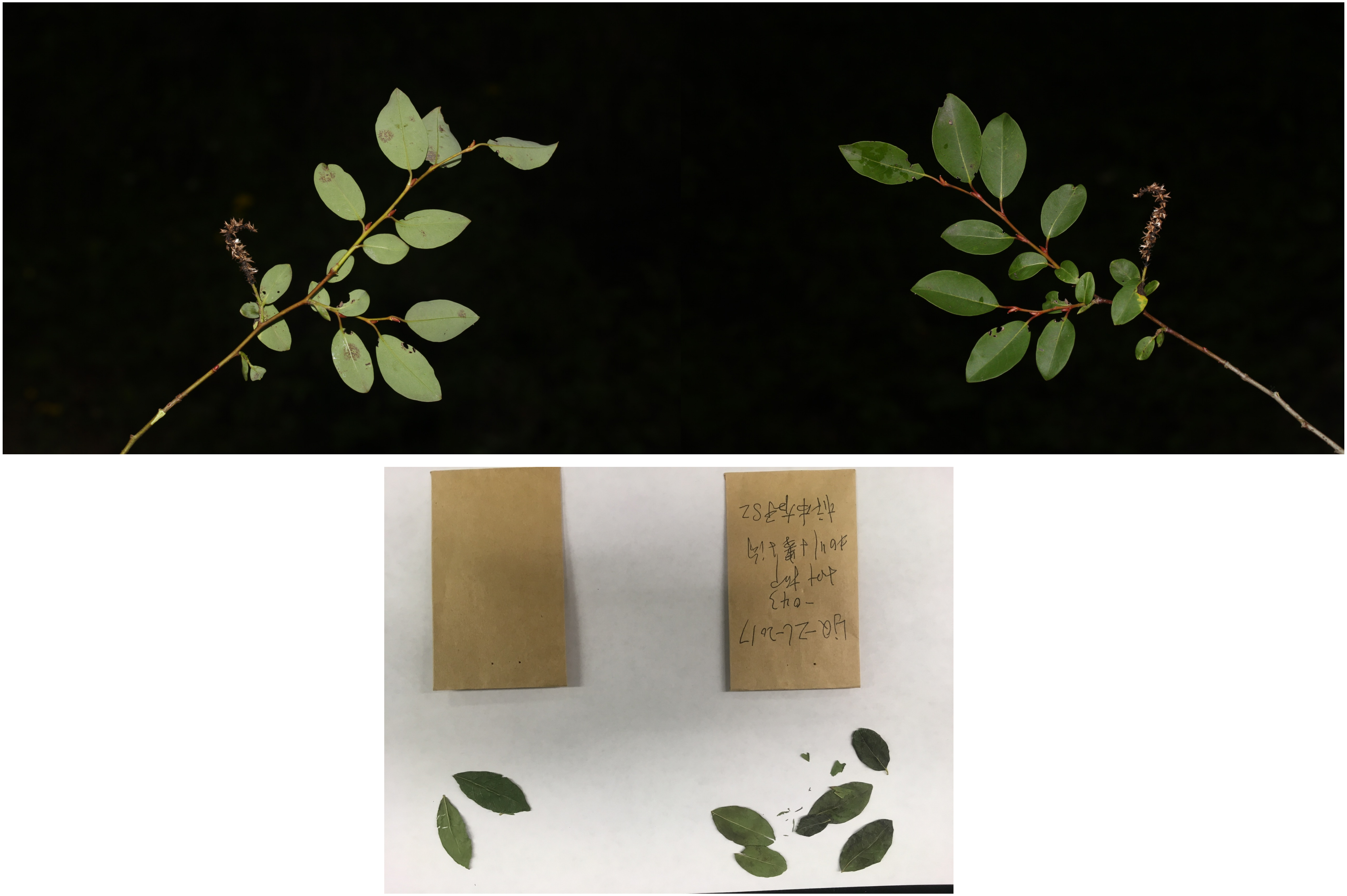

Figure S7. Photographic vouchers for specimens. d. *Salix hypoleuca*

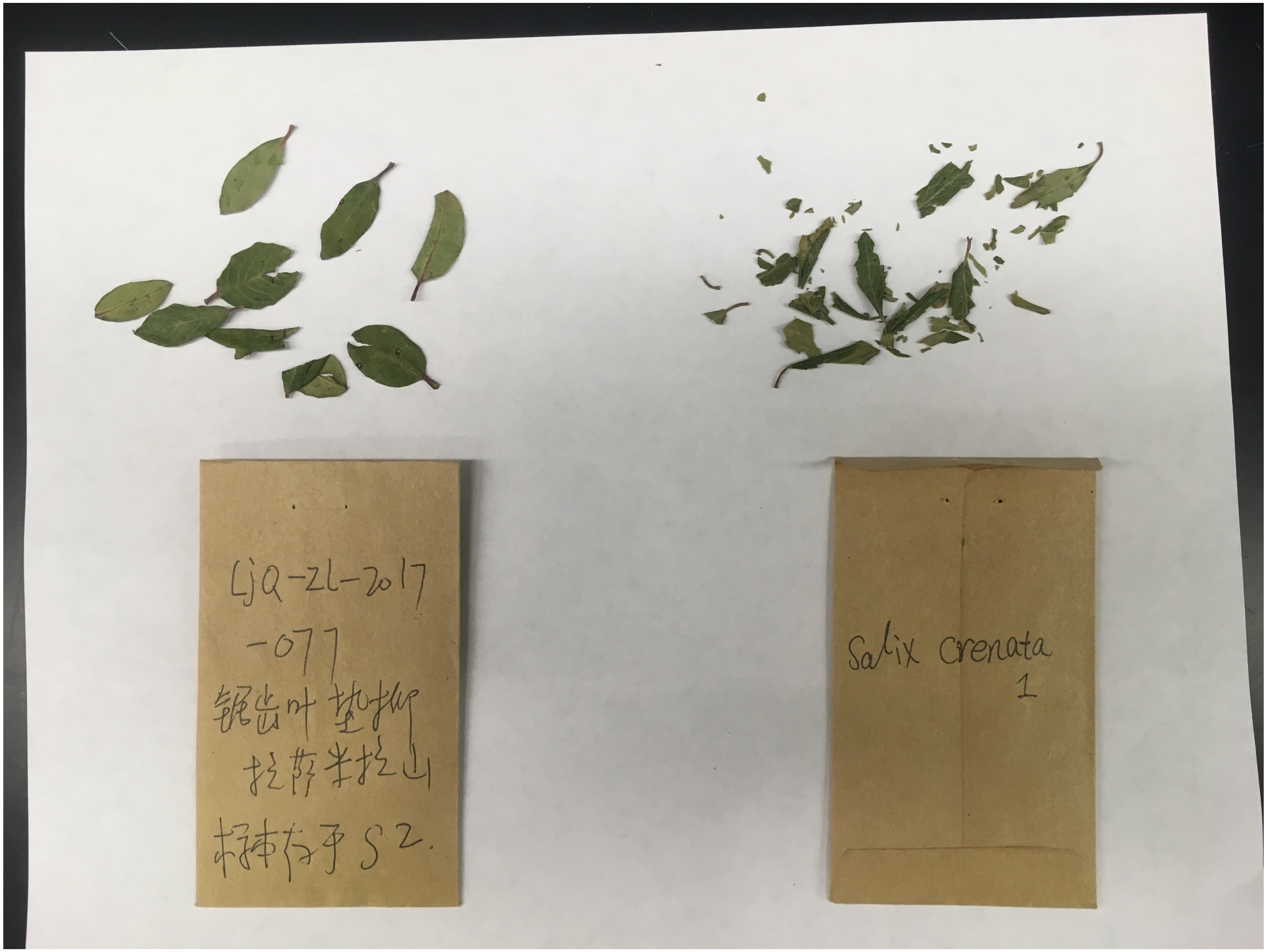


Figure S7. Photographic vouchers for specimens. e. *Salix crenata*

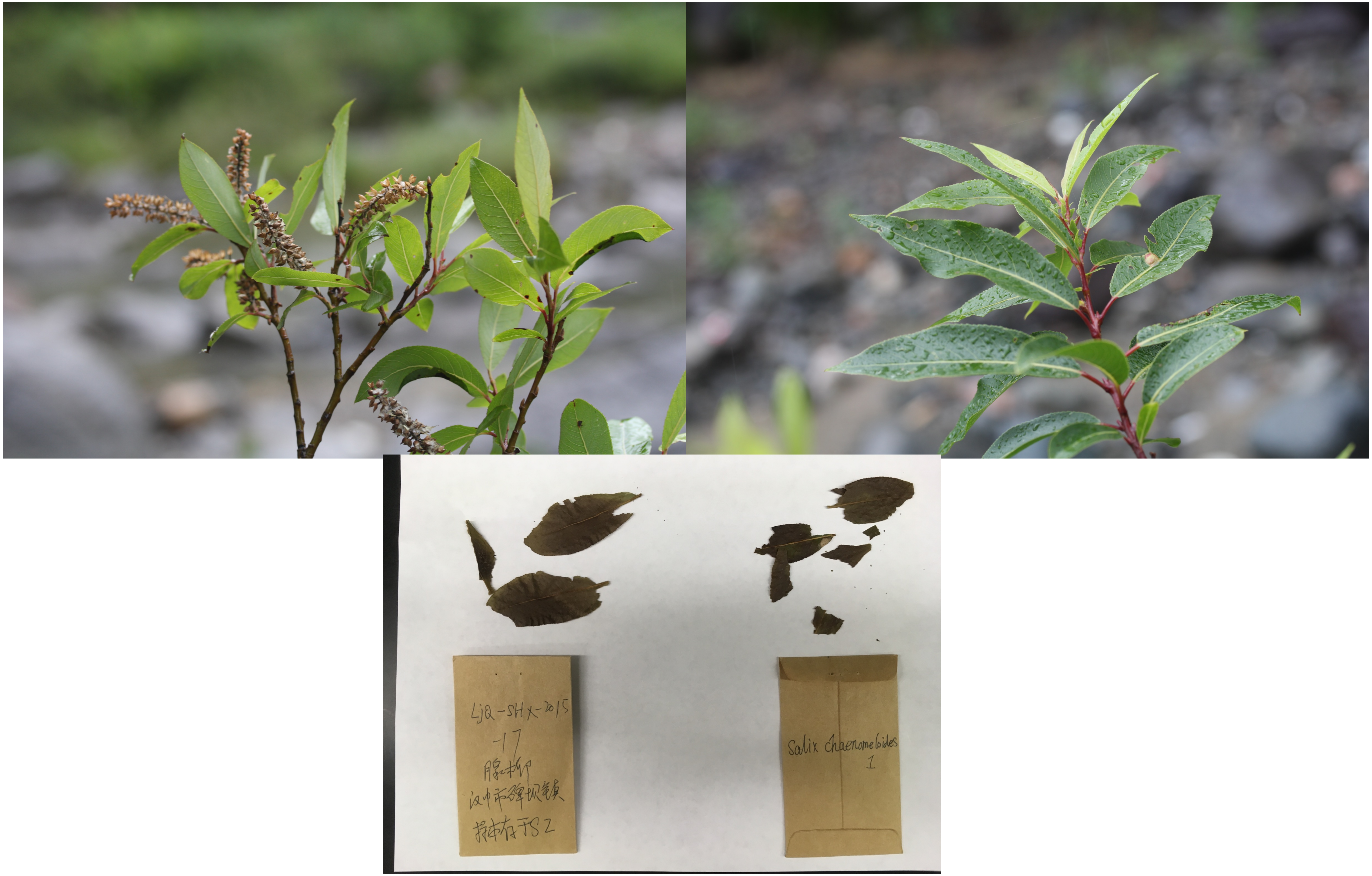


Figure S7. Photographic vouchers for specimens. f. *Salix chaenomeloides*

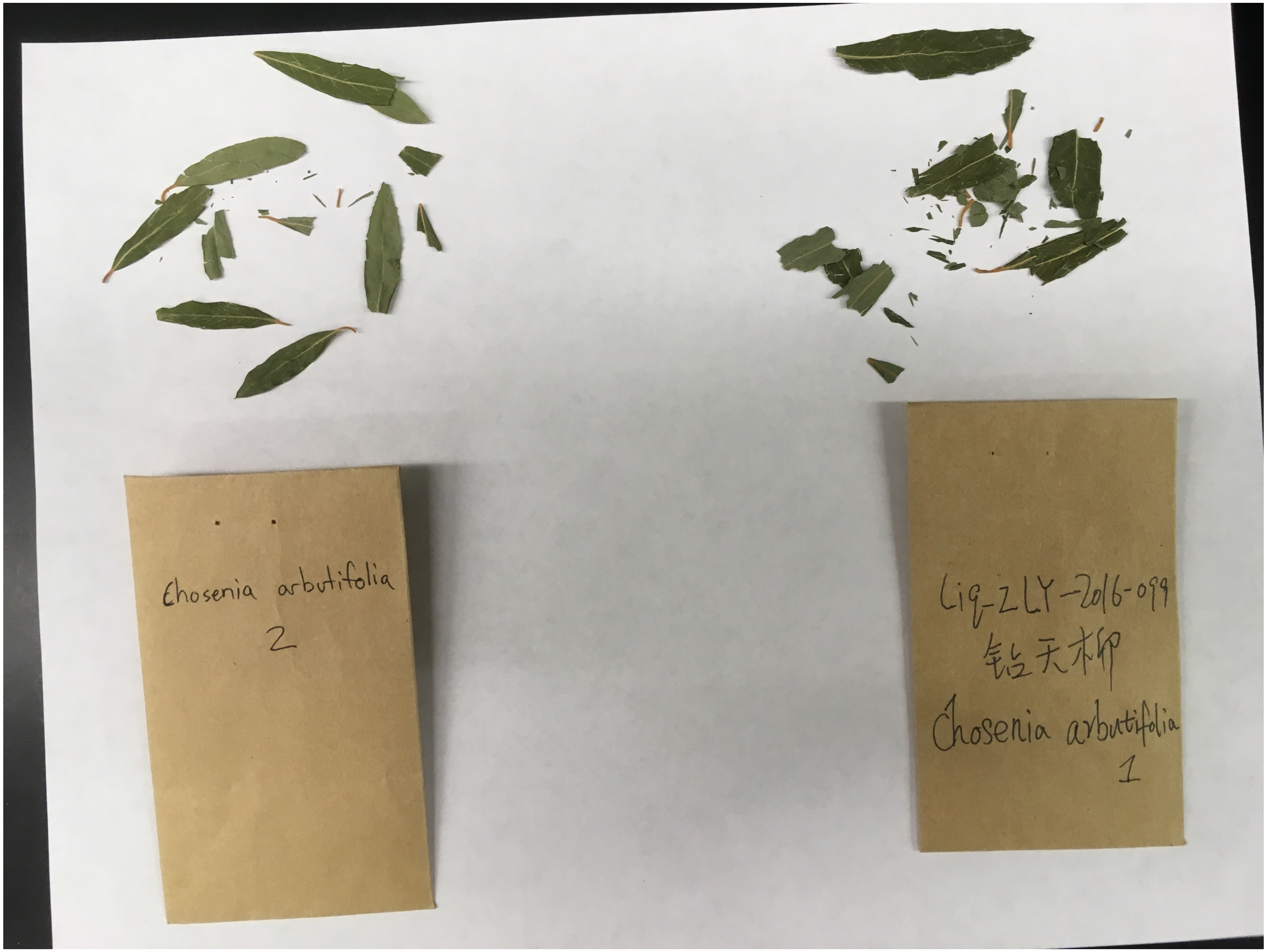


Figure S7. Photographic vouchers for specimens. g. *Salix arbutifolia*
